# Secondary deficiency of neuraminidase 1 contributes to CNS pathology in neurological mucopolysaccharidoses via hypersialylation of brain glycoproteins

**DOI:** 10.1101/2024.04.26.587986

**Authors:** TianMeng Xu, Rachel Heon-Roberts, Travis Moore, Patricia Dubot, Xuefang Pan, Tianlin Guo, Christopher W. Cairo, Rebecca Holley, Brian Bigger, Thomas M. Durcan, Thierry Levade, Jerôme Ausseil, Bénédicte Amilhon, Alexei Gorelik, Bhushan Nagar, Luisa Sturiale, Angelo Palmigiano, Iris Röckle, Hauke Thiesler, Herbert Hildebrandt, Domenico Garozzo, Alexey V. Pshezhetsky

**Author notes:** Correspondence and request for materials should be addressed to: AVP, Sainte-Justine University Hospital Research Center, 3175 Cote Ste-Catherine, Montréal, PQ, H3T 1C5, Canada. Equally contributed as the first authors.

## Abstract

Mucopolysaccharidoses (MPS) are lysosomal storage diseases caused by defects in catabolism of glycosaminoglycans. MPS I, II, III and VII are associated with lysosomal accumulation of heparan sulphate and manifest with neurological deterioration. Most of these neurological MPS currently lack effective treatments. Here, we report that, compared to controls, neuraminidase 1 (NEU1) activity is drastically reduced in brain tissues of neurological MPS patients and in mouse models of MPS I, II, IIIA, IIIB and IIIC, but not of other neurological lysosomal disorders not presenting with heparan sulphate storage. We further show that accumulated heparan sulphate disrupts the lysosomal multienzyme complex of NEU1 with cathepsin A (CTSA), β-galactosidase (GLB1) and glucosamine-6-sulfate sulfatase (GALNS) necessary to maintain enzyme activity, and that NEU1 deficiency is linked to partial deficiencies of GLB1 and GALNS in cortical tissues and iPSC-derived cortical neurons of neurological MPS patients. Increased sialylation of N-linked glycans in brain samples of human MPS III patients and MPS IIIC mice implicated insufficient processing of brain N-linked sialylated glycans, except for polysialic acid, which was reduced in the brains of MPS IIIC mice. Correction of NEU1 activity in MPS IIIC mice by lentiviral gene transfer ameliorated previously identified hallmarks of the disease, including memory impairment, behavioural traits, and reduced levels of the excitatory synapse markers VGLUT1 and PSD95. Overexpression of NEU1 also restored levels of VGLUT1-/PSD95-positive puncta in cortical neurons derived from iPSC of an MPS IIIA patient. Together, our data demonstrate that heparan sulphate-induced secondary NEU1 deficiency and aberrant sialylation of glycoproteins implicated in synaptogenesis, memory, and behaviour constitute a novel pathological pathway in neurological MPS spectrum crucially contributing to CNS pathology.

**Graphical abstract:** 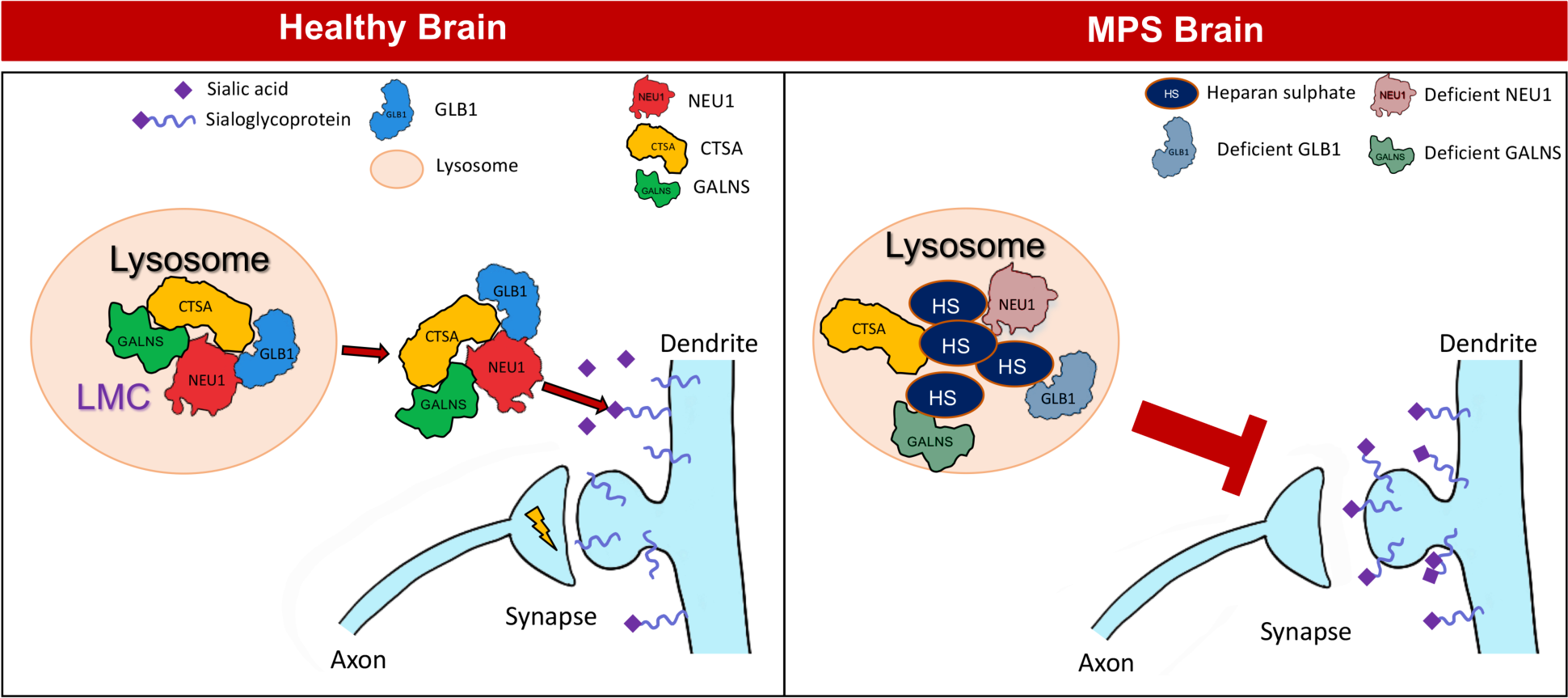

## Introduction

The mucopolysaccharidoses (MPS) are a family of eleven lysosomal storage disorders (LSDs) caused by mutations affecting enzymes involved in degradation of glycosaminoglycans (GAGs) (reviewed in ref. [1]). Neurological MPS are characterized by the accumulation of heparan sulphate (HS) caused by a deficiency of α-L-iduronidase (MPS I), iduronate-2-sulfatase (MPS II), iduronate-2-sulfatase (MPS II), N-sulfoglucosamine sulfohydrolase (MPS IIIA), N-acetyl-α-D-glucosaminidase (MPS IIIB), acetyl-CoA:alpha-glucosaminide N-acetyltransferase (HGSNAT) (MPS IIIC), N-acetylglucosamine-6-sulfate sulfatase (MPS IIID), or β-glucuronidase (MPS VII) [2]. The onset of MPS symptoms in humans varies between cases and subtypes, but generally occurs in early childhood, and severely affected patients usually die during the second or third decade of life. Depending on the subtype, in addition to the central nervous system (CNS), the disease can also progressively affect the peripheral organs and/or skeleton.

Neurological MPS patients experience a range of symptoms including progressive cognitive impairment, sleep disturbance and behavioural problems [1]. Accumulation of HS within neurons and glia significantly affects cell health, triggering secondary pathological cascades including an autophagy blockage and accumulation of misfolded aggregated proteins such as α-synuclein, microtubule-associated protein tau and amyloid-β [3–7]. Other pathological cascades, include synaptic defects, neuroinflammation, astro- and microgliosis [7–11]. In turn, induction of lysosomal biogenesis leads to the accumulation of lysosomes and the increase in the levels and activity of other lysosomal enzymes [4,9,12,13].

In the present study, we report a novel pathological pathway manifesting in neurological MPS spectrum. We show that in contrast to the majority of lysosomal enzymes, the lysosomal neuraminidase 1 (NEU1), an enzyme involved in the cleavage of sialic acids from glycoproteins, shows a remarkable deficiency in the brain tissues of neurological MPS patients and mouse models of the diseases. We further show that NEU1 deficiency in two models of MPS IIIC, knockout, *Hgsnat-Geo* mouse [4] and more severely affected knock-in *Hgsnat^P304L^* mouse [14] is caused by HS-mediated dissociation of its complex with other lysosomal enzymes, cathepsin A (CTSA), β-galactosidase (GLB1) and glucosamine-6-sulfate sulfatase (GALNS), known as lysosomal multienzyme complex (LMC). Although NEU1 and the LMC are mainly known as lysosomal proteins, they also exist on the cell surface and act on cell surface glycoproteins (reviewed in ref. [15]). Our current results show that the secondary NEU1 deficiency in neurological MPS diseases leads to a widespread increase in sialylation of N-linked glycans and synaptic changes reminiscent of neuropathological findings, that are both rescued by lentiviral-mediated genetic correction of NEU1 activity in the MPS IIIC mouse brain.

## Materials and Methods

### Patients

Ethical approval for research involving human tissues was given by the CHU Ste-Justine Research Ethics Board (Comité d’éthique de la recherche FWA00021692, approval number 2020-2365). The National Institute of Health’s NeuroBioBank provided the cerebral tissues, frozen or fixed with paraformaldehyde, from MPS patients as well as age, ethnicity and sex-matched controls (project 1071, MPS Synapse), along with clinical descriptions and the results of a neuropathological examination.

### Animals

The MPS IIIC knock-in model *Hgsnat^P304L^*, MPS IIIC knockout (KO) model, *Hgsnat-Geo*, Tay-Sachs KO mouse model *hexa^−/−^*and NEU1 KO mouse model *Neu1^−/−^* have been previously described [14,16–18]. The animals were housed in the Animal facilities of CHU Ste-Justine, following the guidelines of the Canadian Council on Animal Care (CCAC). The animals were kept in an enriched environment with a 12 h light/dark cycle, fixed temperature, humidity and continuous access to water and food. Approval for the use of the animals in experimentation was granted by the Animal Care and Use Committee of the CHU Ste-Justine (approval number 2023-4090). Frozen brains of previously described mouse models of MPS diseases (MPS I, II, IIIA, IIIB, IVA) [12,19–22], other neurological LSDs (MLD, ML IV, NPC1) [23–25] and their littermate controls were kindly provided by Drs Steven U. Walkley, Volkmar Gieselmann and Shunji Tomatsu. The mice were bred and maintained at the University of Manchester, Albert Einstein College of Medicine, the University of Bönn and the Thomas Jefferson University, Philadelphia. Equal cohorts of male and female mice were studied separately for each experiment and statistical methods used to test whether the progression of the disease, levels of biomarkers or response to therapy were different between male and female animals. Since differences between sexes were not detected, the data for male and female mice and cells were pooled together.

### Real time quantitative PCR

*Neu1*, *Neu3* and *Neu4* mRNA was quantified using a Stratagene Mx3000P QPCR system and mRNA isolated from homogenized half-brains of 2-month-old WT, *Hgsnat^P304L^* and *Hgsnat^Geo^*mice. Briefly, total mRNA was extracted using Trizol (Invitrogen), as described by the manufacturer, and reverse-transcribed using Quantitect reverse transcription kit (Qiagen) as per the manufacturer’s protocol. For the PCR conditions and primer sequences, see Appendix 1.

### Enzyme activity assays

Total acidic neuraminidase, NEU1, β-galactosidase, total β-hexosaminidase and GALNS were measured as previously described [26–28], with minor modifications. Mice were anaesthetised with isoflurane and sacrificed using a CO_2_ chamber. The brain, liver, spleen, kidneys and lungs were extracted and snap-frozen in liquid nitrogen, before being stored at −80 °C. Approximately 50 mg of the tissue were homogenized in 250 µl of water using a sonic dismembrator (Artek Systems Corporation). The NEU1 activity reaction mixture (50 µL) containing 100 µg of total protein, NEU3/NEU4 inhibitor C-9-(4-biphenyl-triazolyl)-DANA (C9-4BPT-DANA; CG17700)[28] at a final concentration of 125 µM, and 4-methylumbelliferone-N-acetyl-neuraminic acid (Sigma-Aldrich) at a final concentration of 250 µM in 25 mM sodium acetate buffer, pH 4.6, was incubated at 37°C for 60 minutes. The reaction was stopped with 950 µl of 0.4 M glycine buffer, pH 10.4. The concentration of the reaction product, 4-methylumbelliferone was measured using a ClarioStar plate reader (BMG Labtech). For all enzymatic assays, blank samples contained all components, except for the homogenate which was added after the termination of the reaction. The β-galactosidase activity reaction mixture containing 10 µl of homogenate, diluted 1:10 in ddH_2_O, 12.5 µl of 0.4 M sodium acetate, 0.2 M sodium chloride buffer, pH 4.2, and 12.5 µl of the fluorogenic substrate 4-methylumbelliferyl β-D-galactoside (Sigma-Aldrich) was incubated at 37°C for 15 min. The reaction was stopped with 965 µl of 0.4 M glycine buffer, pH 10.4 and the product measured as above. The β-hexosaminidase activity reaction mixture containing 2.5 µl of homogenate diluted 1:10, 15 µl of 0.1 M sodium acetate buffer, pH 4.2, and 12.5 µl of 4-methylumbelliferyl N-acetyl-β-D-glucosaminide (Sigma-Aldrich) was incubated for 15 minutes at 37°C. The reaction was stopped with 970 µl of 0.4 M glycine buffer, pH 10.4. The GALNS activity reaction mixture containing 25 µl of the homogenate, 12.5 µl of 1 M sodium acetate, 0.1 M acetic acid, 0.1 M sodium chloride, 5 mM lead acetate, 0.02% sodium azide buffer, pH 4.3, and 25 µl of 4-methylumbelliferyl β-D-galactopyranoside-6-sulphate (Toronto Research Chemicals) was incubated for 3 h at 37°C. Then 12.5 µl of 1.8 M sodium phosphate, 0.02% sodium azide buffer was added, and the incubation continued at 37°C for another hour, when the reaction was stopped with 925 µl of 0.4 M glycine buffer, pH 10.4.

To measure SGSH enzyme activity in cultured iPSC and NPC cells, they were grown in T25 flasks to confluency, washed with 3-5 ml of a cold saline (0.9% sodium chloride) three times, placed on ice and collected with a rubber scraper. SGSH activity was measured as described by Karpova et al. [29], using the synthetic fluorogenic substrate, 4-methylumbelliferyl-2-Deoxy-2-sulfamino-α-D-glucopyranoside Na salt (4MU-α-GLcNs) in a 0.1 M Tris buffer, pH 6.5. Protein concentration was determined using Quick Start™ Bradford Protein Assay (Cat.# 5000201, Bio-Rad Hercules, CA).

### Histochemistry and immunohistochemistry

Sagittal 50 μm-thick sections of cryopreserved mouse brains were prepared as previously described [17]. To analyse bacterial LacZ/β-galactosidase reporter gene expression in the brains of *Neu1^−/−^* mice [17], the sections were washed with 0.1 M phosphate buffer, pH 7.3, supplemented with 2 mM MgCl_2_ and stained with 0.1% (w/v) 5-bromo-4-chloro-3-indolyl-β-D-galactopyranoside (X-GAL) solution in 0.1 M phosphate buffer (pH 7.3) supplemented with 2 mM MgCl_2_, 5 mM potassium ferrocyanide (K_4_Fe(CN)_6_-3H_2_0, Sigma-Aldrich cat. # P-9287) and 5 mM potassium ferricyanide (K_3_Fe(CN)_6,_ Sigma-Aldrich cat. #P-8131) at 37°C overnight. The sections were further washed twice with 0.1 M phosphate buffer, pH 7.3 supplemented with 2 mM MgCl_2_, mounted on the glass slides and images were acquired using a slide scanner (Axioscan 7, Zeiss).

For immunofluorescence, the brain sections were permeabilised and blocked using 0.3% (v/v) Triton X-100, 5% (w/v) bovine serum albumin solution in phosphate-buffered saline (PBS), and stained with primary antibodies diluted in 1% bovine serum albumin, 0.3% Triton X-100 and PBS, overnight at 4°C. This was followed by incubation with an appropriate Alexa Fluor-conjugated secondary antibodies (Life Technologies, see Table 1 for antibodies and dilutions). The slides were mounted using Prolong Gold Antifade Reagent with DAPI (Thermo Fisher Scientific) and images were acquired using a Leica TCS SPE inverted confocal microscope with a 63X oil objective and the same laser settings, exposure time and the number of Z-tacks across animals. Images for all groups of animals were acquired on the same day in the blind fashion.

**Table 1.**
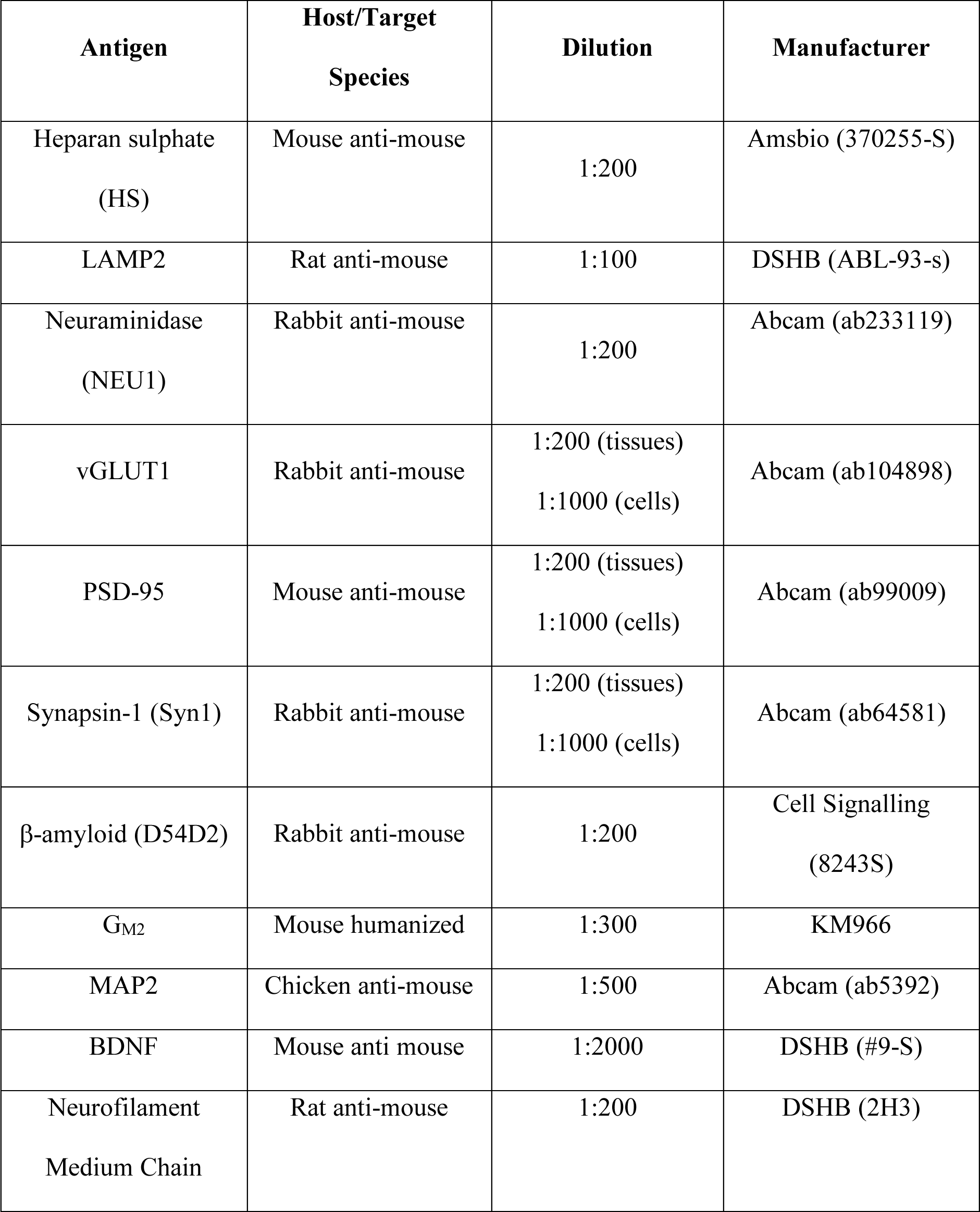
Antibodies and their dilutions used for immunocytochemistry and immunohistochemistry.

| **Antigen** | **Host/Target Species** | **Dilution** | **Manufacturer** |
| --- | --- | --- | --- |
| Heparan sulphate (HS) | Mouse anti-mouse | 1:200 | Amsbio (370255-S) |
| LAMP2 | Rat anti-mouse | 1:100 | DSHB (ABL-93-s) |
| Neuraminidase (NEU1) | Rabbit anti-mouse | 1:200 | Abcam (ab233119) |
| vGLUT1 | Rabbit anti-mouse | 1:200 (tissues)<br>1:1000 (cells) | Abcam (ab104898) |
| PSD-95 | Mouse anti-mouse | 1:200 (tissues)<br>1:1000 (cells) | Abcam (ab99009) |
| Synapsin-1 (Syn1) | Rabbit anti-mouse | 1:200 (tissues)<br>1:1000 (cells) | Abcam (ab64581) |
| β-amyloid (D54D2) | Rabbit anti-mouse | 1:200 | Cell Signalling (8243S) |
| GM2 | Mouse humanized | 1:300 | KM966 |
| MAP2 | Chicken anti-mouse | 1:500 | Abcam (ab5392) |
| BDNF | Mouse anti mouse | 1:2000 | DSHB (#9-S) |
| Neurofilament Medium Chain | Rat anti-mouse | 1:200 | DSHB (2H3) |

Regions of interest were selected from the same cortical areas in the layers IV and V of the somatosensory cortex at the same distance from the dorsal hippocampus. For mice injected with LV, four regions of interest were selected, one from the same area in the dentate gyrus and three from the same areas of CA1. The percentage of stained area was quantified using the ImageJ software for immunohistochemical analysis. The images were z-stacked (z-project, max projection) and converted to B/W (8-bit) format using the same threshold. The percentages of stained areas were averaged within the same animal and compared between different animal groups using nested ANOVA. Densities of PSD-95, VGLUT1, and BDNF positive puncta were estimated by manually counting puncta along the axon at 30 µm increments starting from 10 µm away from the soma. The results were expressed as a mean numbers of puncta/10 µm.

### Analysis of N-linked glycans by mass spectrometry

Proteins from homogenized mouse brains were washed with methanol/chloroform mixture to remove lipids and dried under a nitrogen stream as described [30,31]. Following the denaturation, reduction and alkylation of proteins, N-glycans were released by PNGase F treatment, permethylated and analysed by MALDI-TOF MS and MS/MS using a 4800 Proteomic Analyzer instrument (AB Sciex) as described [31].

### Extraction of secreted HS oligosaccharides from MPS IIIC patients’ urine

Secreted HS oligosaccharides were isolated from MPS IIIC patients urine samples, obtained from the families with informed research consent. Thirty mL of urine was adjusted to a pH 5-6 with acetic acid. After 10 min centrifugation at 1050 g, the supernatant was collected and mixed with 600 µl of 5% cetylpyridinium chloride (CPC). The mixture was then incubated overnight at 4°C and centrifuged for 30 min at 1050 g and 4°C to collect the HS-CPC precipitate. The precipitate was washed twice with 12 mL of ethanol saturated with NaCl; each washing step was followed by 10 min centrifugation at 1050 g and 4°C. After removing the supernatant, 12 mL of 100% ethanol was added to the samples. Following centrifugation, the precipitate was allowed to dry before adding 3 mL of diethyl ether. The supernatant was removed, and the residue dried for another 30 min by a flow of N_2_. Six mL of 0.6 M NaCl was then added to the precipitate, and the mixture was incubated for 3 h at 4°C. After incubation, the samples were cleared by centrifugation and mixed with 24 mL of 100% ethanol. After overnight incubation at 4°C, all steps were repeated and the precipitate (4-5 mg of purified HS oligomers) resuspended in water, freeze-dried and stored at −20°C.

### Analysis of HS treatment on NEU1 activity in cultured bone marrow-derived macrophages (BMDM)

Bone marrow cells (BMC) were collected from the tibia, femur, and iliac bones of WT and *Hgsnat^P304L^* mice. After washing with 70% ethanol and ice-cold PBS containing 1% penicillin and streptomycin, the bones were flushed with ice-cold Dulbecco’s Modified Eagle Medium (DMEM) at both sides to extract bone marrow cells. The cells were collected by centrifugation (10 min at 450 g and 4°C), and resuspended in a red blood cell lysis buffer (155 mM NH_4_Cl, 12 mM NaHCO_3_, and 0.1 M EDTA). After incubation for 30 sec, ice-cold complete DMEM was added to stop the lysis, BMC were collected by centrifugation as before and filtered using a 40 µm Nylon cell strainer. The cells were then cultured in 10 cm Petri dishes in DMEM supplemented with 10% fetal bovine serum (FBS) and containing granulocyte-stimulating factor, obtained from cultured L929 cells [32]. To study the effect of HS, BMDM were cultured for seven days in six-well plates at a density of 5 x 10^6^ cells/well in the absence or presence of HS added in the concentration of 50, 100 and 300 of µg/mL after seeding and after three days in culture, when the medium was replaced. After seven days, the cells were washed with ice-cold PBS and harvested. The total neuraminidase and NEU1 activity were measured in cell homogenates as described above.

### Analysis of brain protein extracts by size-exclusion chromatography

Mouse brains (∼1 g of tissue) were homogenized using a Polytron in 1 ml of 20 mM sodium acetate buffer containing 0.15 M NaCl and 1% Zwittergent 3-14 detergent (Millipore-Sigma 693017), pH 4.75, and cleared by centrifugation for 1 h at 100,000 *g*. A 0.5 ml aliquot of the supernatant was applied to a FPLC Superose 6 column (Pharmacia) and eluted with the same buffer containing 0.1% of Zwittergent at a flow rate of 0.4 ml/min. Thirty 0.5 ml fractions were collected and analysed for neuraminidase, NEU1 and β-galactosidase activities. The molecular masses of the eluted proteins were determined using the calibration curve obtained with the following MW standards (Pharmacia): blue dextran (∼2,000 kDa), thyroglobulin (669 kDa), ferritin (440 kDa), catalase (232 kDa), aldolase (158 kDa), BSA (69 kDa), chymotrypsinogen (25 kDa), and ribonuclease A (13.7 kDa).

### In vitro analysis of HS effects on NEU1 enzymatic activity and solubility

Human NEU1, GLB1 and CTSA were expressed as secreted proteins in Sf9 insect cells, infected with recombinant baculovirus, and purified as previously described [33,34], except that a TEV protease cleavage site was included between the hexahistidine tag and the catalytic domains of NEU1 and CTSA proteins. NEU1 enzymatic activity assays were carried out in 50 mM sodium acetate buffer, pH 4.5, with 100 mM NaCl and 1 mM of the fluorogenic substrate 4-methylumbelliferyl N-acetyl-α-D-neuraminic acid (4MU-NANA, BioSynth EM05195). 20 µM (approximately 1 mg/mL) of TEV-cleaved CTSA was pre-incubated with varying concentrations of HS (sodium salt, from bovine kidney, Sigma H7640) for 30 min at 22 °C, followed by addition of 20 nM of TEV-cleaved NEU1 and incubation for 30 min. After addition of the substrate and incubation for 30 min at 37 °C, the reaction was stopped with 200 mM glycine, pH 10, and the fluorescence of the product 4-methylumbelliferone measured as described above.

To analyze HS effect on the protein solubility, purified recombinant human CTSA, GLB1 or NEU1 were incubated at equimolar amounts (1, 1.41 or 0.79 mg/mL respectively) in the presence of HS (Selleck Chemicals S5992, final concentration 1 mg/mL) in 25 mM sodium acetate buffer, pH 4.5, with 100 mM NaCl for 30 min at 22 °C. Samples were, then, centrifuged at 21000 g for 5 min and pellets solubilized in 8 M urea. Supernatants and pellets were analyzed by SDS-PAGE. CTSA is partially processed into a 30 kDa large and 18 kDa small protein chains by trace amount of endogenous Sf9 proteases during the purification.

### Molecular docking

Coordinates for NEU1 were from the PDB (8DU5)[33]. Initial coordinates for HS tetramers were generated from previous reports [35]. Autodock Vina (version 1.2.5) was used to generate and score docked poses [36], with input files generated in Webina [37]. Poses of heparan sulfate tetramers were then inspected and figures were generated in ChimeraX [38].

### Analyses of polySia-NCAM

Frozen brain samples comprising the right anterior cerebral cortex of 4-month-old WT*, Neu1^−/−^*, *Hgsnat^P304L^*, and *Hgsnat-Geo* mice with average weights of 15.7 mg were homogenized in 30 µl ice cold lysis buffer (100 mM Tris-HCl, pH 7.4, 2 mM EDTA, 1% NP40, 1 mM phenylmethylsulfonyl fluoride, 14 µg/ml aprotinin) per mg sample weight. Lysates were clarified by centrifugation (15 min, 15.700 rcf, 4°C) and total protein concentrations were determined using the Bio-Rad protein assay. Levels of polysialylated neural cell adhesion molecule (polySia-NCAM) were determined by a sandwich ELISA [39] adapted to mouse samples. Briefly, 96 well half area microplates (Cat.# 675101, Greiner Bio-one, Frickenhausen, Germany) were coated with polysialic acid (polySia)-specific monoclonal antibody 735 [40] (mouse IgG_2a_; 5 µg/ml in PBS, 25µl per well) for 2 h at room temperature (RT), washed with 0.1 M phosphate buffer, pH 7.4 containing 0.1% Tween20, and blocked overnight at 4°C with 1% bovine serum albumin (BSA, Merck, Cat# 7906) in PBS (200 µl per well). Ten µl of cortex lysate, diluted to a concentration of 50 µg of protein per ml, was added to each well, and the plates were incubated overnight at 4°C. After washing, wells were incubated with the NCAM specific monoclonal antibody OB11 (Merck, Cat.# C9672, mouse IgG_1_; 1:1000), followed by horseradish peroxidase (HRP)-conjugated anti-mouse IgG_1_ (Jackson Cat.# 115-035-205; 40 ng/ml), both for 1h at RT with 10 µl per well in blocking buffer). HRP substrate 3,3’,5,5’-Tetramethylbenzidin (TMB, Merck, Cat.# T2885; 0.1 mg/ml in citrate buffer, pH 4.9, with 0.001% H_2_O_2_) was added, the reaction was stopped after 25 min with 1 M H_2_SO_4_ (10 µl per well), and absorbances were measured at 450 nm against 550 nm as a reference wavelength. Each sample was analyzed by four independent experiments and values were normalized to the mean value of the WT samples. Sample treated with endosialidase (6 µg/ml, 40 min at 37°C), degrading polySia with high specificity [41], were used as negative ELISA controls. Endosialidase treatment reduced absorption levels to <1% of the values for untreated samples. Western blot analysis of cortex lysates was performed with polySia-specific mAb 735 and NCAM-specific mAb H28 as described previously [42]. Detection and densitometric quantification were performed with the Odyssey Infrared Imaging System and Image studio 4.0.21 software (LI-COR Biosciences, Bad Homburg, Germany).

### Generation and maintenance of induced pluripotent stem cells (iPSCs)

The induced pluripotent stem cells (iPSCs) were generated from the fibroblasts of an MPS IIIA patient, obtained from the Coriell Institute for Medical Research, NJ, USA (line GM01881), and a healthy sex and age-matched control from the biobank of CHUSJ. The generation and use of human iPSCs was approved by the Research Ethics Committee of CHUSJ (approval number 2022-3817). iPSCs were tested for mycoplasma and reprogrammed at CHUSJ Cell reprogramming and genome editing core using the CytoTune™-iPS 2.0 Sendai Kit (Life Technologies) [43]. Once generated, the iPSCs lines were maintained in a 5% CO2, 5% O_2_, 37 °C incubator using Matrigel-coated dishes and mTeSR™ Plus (StemCell) media as described [44]. The expression of pluripotency markers Oct4, SSEA4, TRA-160, and SOX2 in all iPSCs lines was confirmed by immunofluorescence.

### Generation of cultured cortical neuronal progenitor cells and cultured cortical iPSC-derived neurons

After two passages, iPSCs were differentiated into a monolayer of forebrain committed neural progenitor cells (NPCs) by dual SMAD inhibition, with FGF-8 used instead of FGFb-2 [45]. NPC were cultured using a neuronal induction media (DMEM/F12) for 3 weeks and analyzed by immunofluorescence to confirm the expression of the endoderm markers, Nestin and PAX6, as well as the neuronal markers, NeuN, axonal β-tubulin III (clone TUJ1) and Syn1. After three weeks in culture, NPC were further differentiated into cortical-specific neurons as described [46] but MPS IIIA patient cell lines were transduced with the LV-CTSA-IRES-NEU1-GFP or control LV-GFP virus at a multiplicity of infection of 10 (MOI 10) for 24 h before the switch to the differentiation media. Neurons were then cultured to day 28 to achieve a complete differentiation and maturation. All cells expressed fidelity markers Nestin/Pax6/Tuj1/Syn1) as well as the cortical specific marker T-box brain 1 (TBR1). Fluorescence activated cell sorting (FACS) analysis confirmed that >80% of cells were TBR1-positive and ∼72% of cells were NeuN/TBR1-positive, demonstrating a high degree of conversion. The primary enzymatic deficiency of SGSH in MPS IIIA cells was confirmed at both iPSC and NPC stages. Size and abundance of LAMP2-positive perinuclear puncta, a marker for lysosomal storage/increased lysosomal biogenesis phenotype [47], were measured by immunocytochemistry at the NPC stage.

### Stereotaxic injections of mice with lentiviral vectors

The previously described [48] CTSA-IRES-NEU1-GFP biscistronic HIV-1-based recombinant vesicular stomatitis virus glycoprotein-pseudotyped (VSVg) LV and control GFP-LV were generously provided by Dr. Jeffrey A Medin (Medical College of Wisconsin, Milwaukee, USA). LV titers were 1.4 × 10^8^ TU/ml for CTSA-IRES-NEU1-GFP LV and 8.3 × 10^8^ TU/ml for control NEU1-GFP LV. Postnatal days 18 and 19 (P18, P19) mice were anesthetized with 5% isoflurane and oxygen in an induction chamber. During the surgery, mice remained on a heating pad to prevent hypothermia and the mouse’s head was secured in an induction cone with a flow of 2 L of oxygen containing 2% isoflurane per min. Mice were further immobilized by fixing their heads with ear bars and injected subcutaneously with a non-steroidal anti-inflammatory drug Carprofen/Rimadyl (Zoetis, DIN 02255693, 0.1 mL of Carprofen mixed with 9.9 mL of 0.9% Sodium Chloride, 0.15 mL per mouse) and a lubricant OptixCare was applied to protect the eyes. After removal of the fur on top of the head and disinfecting the surgical area with 10% povidone-iodine solution (Laboratoire Atlas) the skin was cut open. One hole was made on each side of the brain using a microdrill (Foredom, K.1070 Micromotor kit). The coordinates were 1.5 mm on each hemisphere’s M/L axis and 2.2 mm on the A/P axis. The holes were less than 1 mm in diameter and deep enough to pierce the skull without damaging the dura mater. To remove bone fragments around the holes, a saline solution was used to flush the surrounding areas.

The glass capillary needles for injections were prepared using glass capillaries (3.5” Drummond #3-000-203-G/X) and a micropipette puller (Sutter Instrument Company). After backfilling the glass capillary needles with mineral oil (Sigma BioReagent, #MKCM5718), the needles were fixed onto the Nanoliter Injector (Drummond Scientific Company, Nanoject III #3-000-207) and the Nanoliter micropipette was moved to the injection site (1.5 mm M/L, 2.2 mm A/P). For each hemisphere, the micropipette was first lowered to a depth of 2.2 mm to target the hippocampus, and then, to 1.2 mm to target layers IV and V of the cortex. The LV was injected at a speed of 2 nL/second using the Nanoliter injector, and each injection lasted about seven minutes. After injecting 900 nL of the virus into the hippocampus, we waited six minutes for the viral solution to set down before moving the needle to the cortex. After all the injections were completed, the wound was sutured with ethilon nylon suture (Ethicon, PMP346), and an antibiotic cream (Polysporin, Jonhson & Jonhson inc, DIN 02237227), was applied to the surgical zone. Finally, another dose of diluted Carprofen (0.15 mL) was injected subcutaneously into the mouse, which was then removed from the stereotaxic frame and returned to its cage. All the injected mice were put on a wet diet, and their condition was observed daily following the surgery.

### Behavioural analysis

Novel object recognition (NOR) test was used to measure short-term memory. One day before the experiment, mice were habituated to a white plexiglass box (45 cm length x 45 cm width x 40 cm height) for 10 min and, then, returned to their home cages. The next day, mice were placed individually in the testing chamber facing the wall at the opposite side of two identical blue plastic cylinders measuring 6.5 x 7.5 cm and placed at a 15 cm distance from the walls. The animals were allowed to explore the objects for 10 min before returning to their home cages. One hour later, the mouse was placed back into the testing chamber, which contained one of the original and a new object, a red plastic cube measuring 6.5 x 6.5 x 6.5 cm. Each trial with the novel object lasted for 10 min. Between each session, the testing chambers and the objects were washed with 70% ethanol to eliminate any olfactory cue bias. Each session was video-recorded and analyzed manually by an operator blinded to the animal genotype and treatment. The exploration time was counted when the head of the mouse was within a 3 cm radius of the object, and when the animal was looking at the object, sniffing the object, or touching the object with its snout. Exploration time was not counted when the mouse was within the exploration zone but not actively exploring the objects, or if the animal was grooming itself or sitting on top of the toys. The discrimination index (DI) was calculated as the difference in time spent exploring the new and original objects divided by the total exploration time.

Open field (OF) test was used to measure anxiety and activity as described [4]. Mice were habituated to the testing room for 30 min before the experiment, placed in the center of an open-field arena (45 cm length x 45 cm width x 40 cm height) under dimed light conditions (∼30 lux) and allowed to explore it for 20 min. Each session was recorded and analyzed by the Smart 3.0 software. The following parameters were measured: the number of entries in the center, the total distance traveled, the percent of time spent in the center zone, and the distance traveled in the center zone. The arena was cleaned with 70% ethanol between the trials. OFT was always performed one hour into the mouse light cycle by the same investigator (TM.X.).

### Statistical analysis

Statistical analyses were performed using Prism GraphPad software (GraphPad SoftwareSan Diego, CA). All data were analyzed for normal distribution using the Shapiro-Wilk normality test. Significance of the difference was determined using t-test (normal distribution) or Mann-Whitney test when comparing two groups. One-way ANOVA, nested ANOVA (normal distribution) or Kruskal-Wallis test followed by Dunn or Tukey multiple comparisons tests were used when comparing more than two groups. Two-way ANOVA was used for two-factor analysis. A P-value of 0.05 or less was considered significant.

## Results

### Secondary deficiency of NEU1 in tissues of neurological MPS patients and mouse models

To characterize the biomarkers of CNS pathology in neurological MPS spectrum, we analysed frozen somatosensory cortex post-mortem tissues of one MPS I, one MPS II, two MPS IIIA, one MPS IIIC, and two MPS IIID patients and 7 controls matched for age, gender and ethnicity (project 1071, MPS Synapse, see [7] and Supplementary Table S1 for patient description). The combined activities of lysosomal β-hexosaminidases A and B were significantly increased in all patient samples except for MPS II where they showed a trend towards an increase (Figure 1A). This was consistent with the previously reported increase of lysosomal biogenesis and TFEB-driven expression of lysosomal genes in tissues and cells of patients affected with lysosomal storage diseases [4,9,12,13]. The changes were also reminiscent to the previously described increase of lysosomal enzyme activities and lysosome-associated membrane protein 2 (LAMP-2) levels in *Hgsnat^P304L^* and *Hgsnat-Geo* mouse models of MPS IIIC [14,16]. Unexpectedly, the total acidic neuraminidase activity, representing combined activities of NEU1, NEU3 and NEU4 isoenzymes, was either reduced or showed a trend towards a reduction (Figure 1B). Furthermore, the activity of NEU1 isoenzyme, measured in the presence of C9-4BPT-DANA, a potent inhibitor of both human and mouse NEU3 and NEU4 (IC_50_ 0.7 μM and 0.5 μM, respectively) with minimal activity against NEU1 [28], showed a drastic reduction in all brain tissues compared to their respective controls (Figure 1C). In contrast, NEU3/NEU4 activity deduced by subtraction of NEU1 from the total neuraminidase activity was unchanged (not shown). This indicates that NEU1 is the neuraminidase affected in the MPS tissues. Similarly, total neuraminidase and NEU1 activities were drastically reduced in the brain, liver, and lungs, but not in the spleen and the kidney of both *Hgsnat^P304L^* and *Hgsnat-Geo* MPS IIIC mouse models compared to wild type (WT) controls (Figure 1D and 1E), while the total β-hexosaminidase and β-galactosidase activities were either increased or similar to those of controls (Figure S1A and S1B).

**Figure 1.**
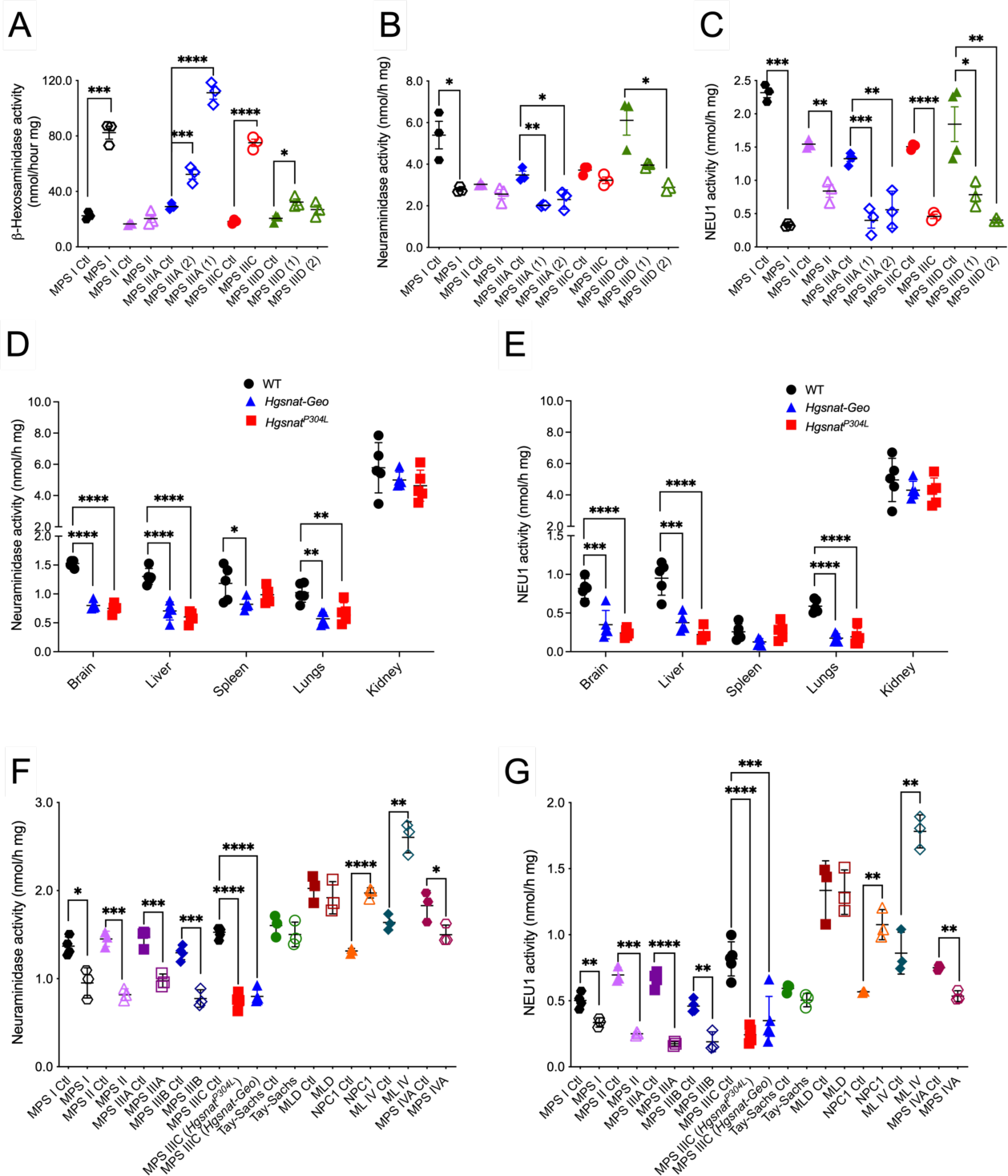
NEU1 activity is reduced in brain samples of neurological MPS patients, brain tissues of neurological MPS mouse models and tissues of MPS IIIC mouse models. **(A-C)** Total β-hexosaminidase activity **(A)**, total acidic neuraminidase activity **(B),** and NEU1 activity in the presence of a specific NEU3/4 inhibitor, C9-4BPT-DANA **(C)** in post-mortem cortical tissues of neurological MPS I, MPS II, MPS IIIA, MPS IIIC, and MPS IIID patients and their corresponding age, sex and ethnicity-matching controls (Ctl, Supplementary Table S1). NEU1 activity is reduced in all MPS patient’s samples compared to those of controls. (**D-G)** Total neuraminidase **(D)** and NEU1 **(E)** activities are reduced in the brain, liver and lungs of MPS IIIC mice compared to WT controls. Total neuraminidase **(F)** and NEU1 **(G**) activities are also reduced in the brains of the mouse models of neurological MPS disorders and MPS IVA compared to respective WT littermates, but are similar to WT or increased in other LSD models. ML IV, mucolipidosis IV; NPC1, Niemann-Pick type C1; MLD, metachromatic leukodystrophy. Graphs show individual values and means (±SD) of 3-4 technical replicates per patient (**A-C**) or 3-5 mice (**D-G**). The patient data represent. *P < 0.05, **P < 0.01, ***P < 0.001, ****P < 0.0001, determined by unpaired t-tests or Mann-Whitney tests.

To test whether the secondary deficiency of NEU1 is shared by other lysosomal disorders, we analysed total neuraminidase and NEU1 activities in the brain tissues of available MPS mouse models (MPS I, II, IIIA, IIIB and IVA) and models of other neurological LSD (MLD, Tay-Sachs, Niemann-Pick type C1, and ML IV). All mouse models of neurological MPS I, II, IIIA, IIIB and IIIC, characterized by HS accumulation exhibited strongly reduced total neuraminidase and NEU1 activity while activity levels in mouse models of other LSD were unaffected (MLD, Tay-Sachs), or even increased (Niemann-Pick type C1, ML IV) compared to their corresponding WT littermate controls (Figure 1F, G). In the brains of the mouse model of MPS IVA, a non-neurological MPS, manifesting with some degree of HS storage in both patients and the mouse model [49,50], the activity of NEU1 was decreased, but only by about 30%. This established NEU1 deficiency as a common phenomenon across all neurological MPS diseases in both mouse models and human patients.

### Secondary deficiency of NEU1 is associated with lysosomal storage of HS

X-Gal staining of brain sections from *Neu1* KO (*Neu1^−/−^*) mice expressing the bacterial LacZ/β-galactosidase reporter gene in the endogenous *Neu1* locus [17] identified the hippocampus, cerebellum, olfactory bulb, paleocortex and deep layers of frontal cortex as the brain areas with the highest *Neu1* expression (Figure S2). These regions were micro-dissected from brains of WT and MPS IIIC mice of both sexes and analysed for total neuraminidase and NEU1 activities, which were found to be reduced in the hippocampus, paleocortex, frontal cortex and cerebellum, but not in the olfactory bulb (Figure 2A-E).

**Figure 2.**
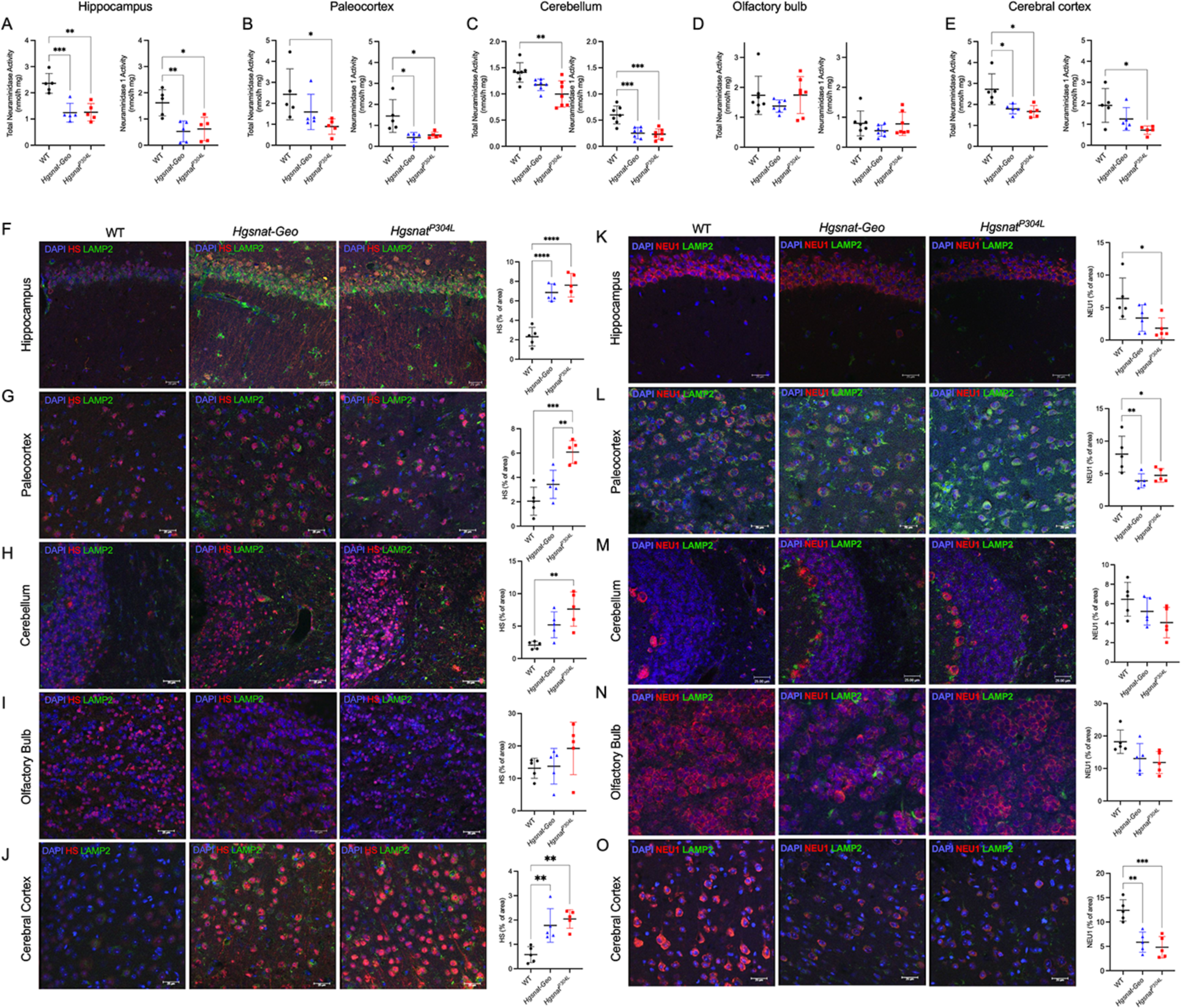
Secondary deficiency of NEU1 in the MPS IIIC mouse brain coincides with HS accumulation. **(A-E)** Total neuraminidase and NEU1 activity in the hippocampus (**A)**, paleocortex (**B)**, cerebellum (**C)**, olfactory bulb (**D)**, and frontal cortex (**E)** of 6-month-old WT, *Hgsnat-Geo,* and *Hgsnat^P304L^* mice. NEU1 activity was reduced in the hippocampus, paleocortex, cerebellum, and cerebral cortex but not in the olfactory bulb. **(F-O)** Representative images and quantifications of HS **(F-J)** and NEU1 **(K-O)** immunoreactivity (red) of brain tissues of 6-month-old WT, *Hgsnat-Geo,* and *Hgsnat^P304L^* mice co-labeled with antibodies against LAMP-2 (green), and DAPI (blue). HS is increased in the hippocampus **(F)**, paleocortex **(G)**, and frontal cortex **(J)** of MPS IIIC compared to WT mice. *Hgsnat^P304L^* but not *Hgsnat-Geo* mice also show an accumulation of HS in the cerebellum **(H)**. The level of HS in the olfactory bulb **(K)** is similar in MPS IIIC and WT mice. NEU1 is decreased in the hippocampus **(K)**, paleocortex **(L)**, and frontal cortex **(O)** of MPS IIIC mice but not in the cerebellum (**M)** and olfactory bulb **(N)**. HS shows a trend for increase and NEU1 for decrease in *Hgsnat^P304L^* compared to *Hgsnat-Geo* mice. Scale bars: 25 µm. Immunoreactivity was evaluated as percentage of total area using ImageJ software. Graphs display individual data, and means (±SD) from 5-7 mice. *P < 0.05, **P < 0.01, ***P < 0.001, ****P < 0.0001, determined by nested one-way ANOVA with Tukey post hoc tests. In all experiments mice of both sexes were used and the data were pooled since no difference was detected between males and females.

To test if the reduction of NEU1 activity was associated with the changes in the levels of NEU1 proteins and HS accumulation, brain sections of 6-month-old WT, *Hgsnat-Geo,* and *Hgsnat^P304L^* mice were analyzed by immunohistochemistry (IHC) with NEU1- and HS-specific antibodies. Levels of NEU1 were decreased and HS was drastically increased in the hippocampus, paleocortex, cerebral cortex and cerebellum of *Hgsnat^P304L^* mice, but not in the olfactory bulb (Figure 2F-O). Changes in *Hgsnat-Geo* mice were similar, but less pronounced. Together, these results suggest that the accumulation of HS in the brain is associated with deficiencies of NEU1 protein and activity.

As described previously, intracerebroventricular (ICV) injections of an AAV “true type” (AAV-TT) vector expressing human WT HGSNAT rescued HGSNAT deficiency in *Hgsnat-Geo* mice, and significantly reduced HS levels (on average by ∼46%) [51]. Concurrently, levels of neuroinflammation were reduced and behavioural deficits in the Y-maze and the Open Field tests ameliorated[51]. We, therefore, used the previously collected samples to test whether the treatment also rescued the secondary NEU1 deficiency. As shown by immunohistochemistry, HS accumulation was reduced in *Hgsnat-Geo* mice with AAV-mediated HGSNAT expression, and this was paralleled by an almost complete restoration of NEU1 protein levels (Figure 3A, B).

**Figure 3.**
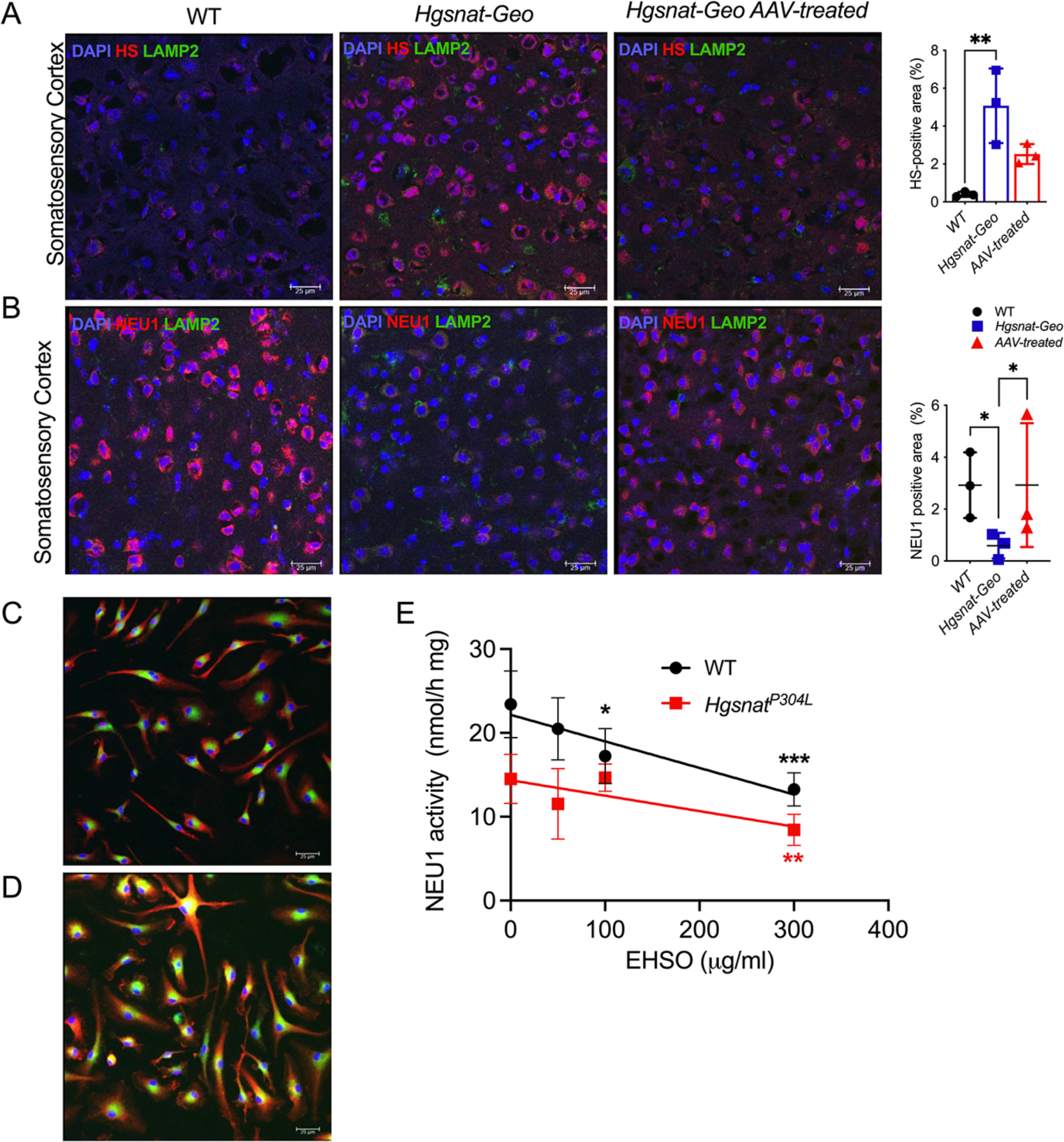
HS accumulation causes deficiency of NEU1. **(A-B)** HS storage is reduced and NEU1 levels increased in MPS IIIC mice treated with intracerebral injections of AAV-TT-HGSNAT vector. Brain sections of 6-month-old WT, untreated *Hgsnat-Geo* mice, and *Hgsnat-Geo* mice treated with AAV-TT-HGSNAT were labeled for **(A)** HS (red) and LAMP2 (green) or **(B)** NEU1 (red) and LAMP2 (green). Nuclei were counterstained with DAPI (blue). HS shows a trend towards a decrease and NEU1 is increased in the brains of AAV-TT-HGSNAT-treated *Hgsnat-Geo*. Panels show representative images of the somatosensory cortex taken with a 40x objective; scale bars: 25 µm. Data on the graphs show individual values, means ± SD, (n=3, five areas per mouse). Significance was determined by nested one-way ANOVA and a Tukey post hoc test. **(C-F)** Administration of exogenous HS oligomers (EHSO) to cultured BMDM causes secondary NEU1 deficiency. HS levels in cultured *Hgsnat^P304L^* BMDM untreated **(C)** or treated with 300 µg per mL of EHSO **(D)** were analysed by IHC. **(E)** NEU1 activity is progressively reduced in the BMDM cultured in the presence of increasing concentrations of EHSO, as indicated. Confocal images were taken using a 40x objective, scale bars: 25 µm. Data on the graphs show individual data, means ± SD, n=4. Significance was determined by two-way ANOVA with a Dunnett post hoc test.

Cultured bone marrow-derived macrophages (BMDM) from WT and *Hgsnat^P304L^*mice were used to determine whether reduced NEU1 activity was a consequence of HS accumulation in lysosomes. As shown before, extracellular HS oligomers (EHSO), purified from the urine of MPS IIIC patients are readily taken up by cultured phagocytic cells such as macrophages or microglia [52]. In our experiments, EHSO treatment increased HS in the lysosomal lumen (Figure 3C, D), and reduced Neu1 activity of WT BMDM in a dose-dependent manner (Figure 3E). Consistent with the results obtained with brain tissue samples (see Figure 1B), NEU1 activity in untreated BMDM from *Hgsnat^P304L^* mice was about 50% of the level in WT cells, but also progressively reduced with increasing concentrations of EHSO (Figure 3E). Notably, when EHSO at similar concentrations were added to the homogenates of mouse kidney tissues during the HGSNAT assay, we did not observe any reduction of the enzymatic activity showing that in this concentration range the compound does directly inhibit NEU1 (Figure S3).

### HS induces secondary deficiency of NEU1 and disruption of the lysosomal multienzyme complex

To test if HS induced the secondary deficiency of NEU1 by down-regulating the *Neu1* gene expression, *Neu1* mRNA levels were measured by quantitative RT-PCR. However, comparable *Neu1* mRNA levels were detected in the brains of WT, *Hgsnat^P304L^* and *Hgsnat-Geo* animals (Figure 4A). This result was also consistent with neuraminidase expression levels, derived from a previously published data set on gene expression in the hippocampus of MPS IIIC and WT mice [14] (Figure 4B). Together with the reduction of NEU1 protein in the brain of *Hgsnat^P304L^*and *Hgsnat-Geo* mice revealed by immunofluorescence analysis (Figure 2K-O), these data suggest that the NEU1 levels are altered at the post-transcriptional level.

**Figure 4.**
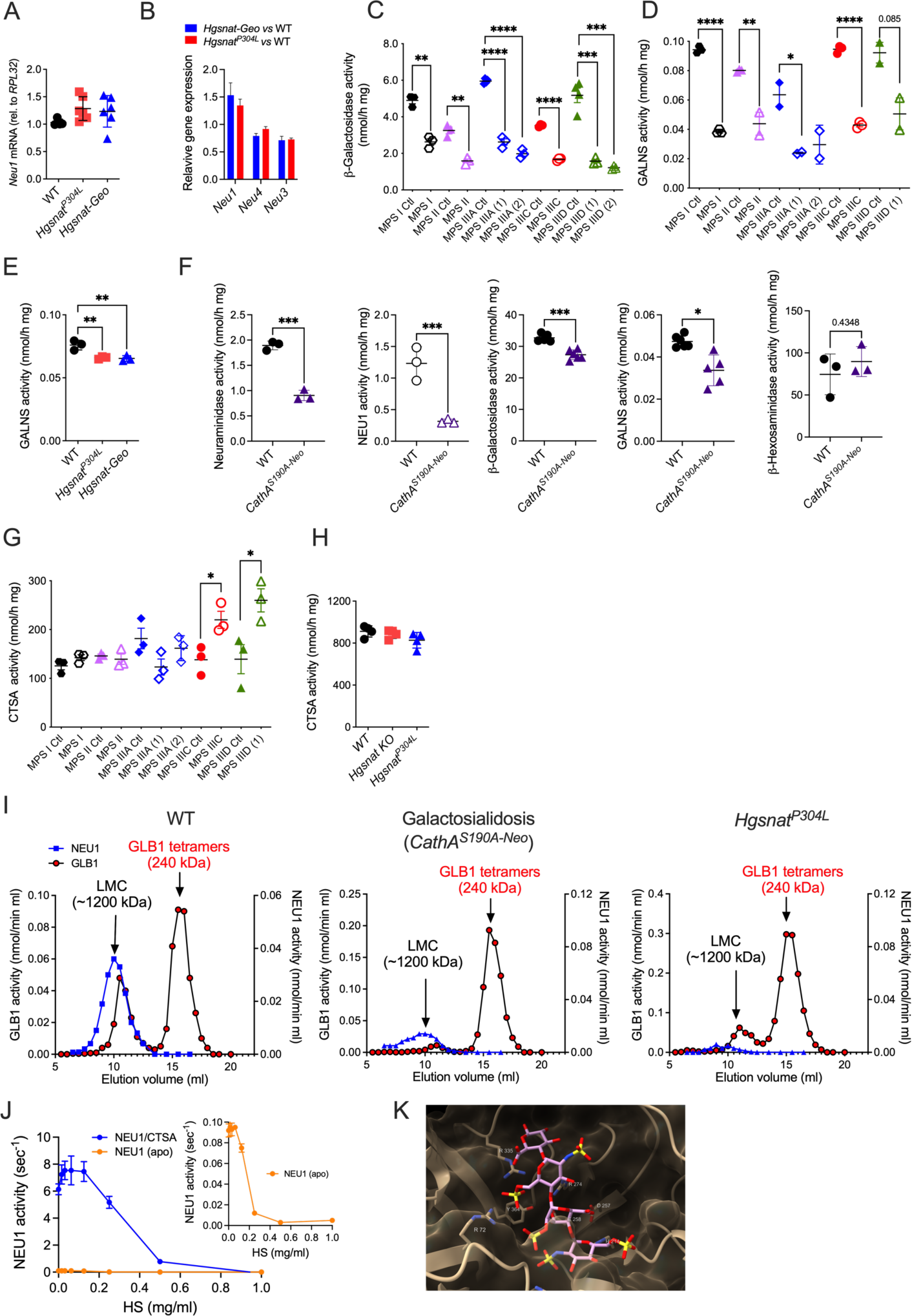
NEU1 deficiency in tissues and cells with lysosomal storage of HS is associated with the disruption of the LMC. **(A, B)** *Neu1* mRNA levels are similar in brain tissues of WT and MPS IIIC mice. **(A)** Neu1 mRNA was measured by real time quantitative PCR using total mRNA extracted from half-brains of 2-month-old WT, *Hgsnat^P304L^* and *Hgsnat-Geo* mice. The values were normalized for the expression of RPL32. Individual results and means (±SD), n=4-5 are shown. **(B)** Relative expression of *Neu1*, *Neu3* and *Neu4* was measured in the hippocampi of WT, *Hgsnat^P304L^* and *Hgsnat-Geo* mice by total mRNA sequencing [14]. The bar graph displays gene expression in *Hgsnat^P304L^* and *Hgsnat-Geo* relative to WT mice. Data show means (±SD), n=3. **(C)** GLB1 activity is reduced in brain samples of neurological MPS patients. Acidic β-galactosidase activity was measured in post-mortem cortical tissues from patients affected with MPS I (561), MPS II (902), MPS IIIA (3617 and 563), MPS IIIC (6194), and MPS IIID (5411 and 5424), and from corresponding age, sex and ethnicity-matching controls. Data show means (±SD) of three technical replicates. ***P < 0.001, ****P < 0.0001, determined by unpaired t-test. **(D)** GALNS activity is reduced in MPS patient brain samples compared to age, sex and ethnicity matched controls. Data show individual results and means (±SD) of three technical replicates. *P < 0.05, **P < 0.001, ****P < 0.0001, determined by unpaired t-test analysis. **(E)** GALNS activity is slightly reduced in brain tissues of 4-month-old MPS IIIC compared to WT mice. Data show individual results and means ±SD, n=3. *P < 0.05, determined by one-way ANOVA with a Tukey post hoc test. **(F)** NEU1, but not GLB1 or GALNS, is deficient in the brain tissues of galactosialidosis *CathA^S190A-neo^* mice. Data show individual results and means (±SD), n=3. **(G-H)** CTSA activity in brain samples of human MPS patients **(G)** and MPS IIIC mice **(H)** is similar or higher than that of corresponding controls. Data show individual results and means (±SD) of three technical replicas for human samples or 3 mice. **(I)** The gel-filtration profiles of total protein extracts from the brain tissues of MPS IIIC *Hgsnat^P304L^*, WT and *CathA^S190A-Neo^* mice. GLB1 and NEU1 activities in the eluted fractions is plotted on the left and right Y-axes, respectively. (**J**) HS inhibits NEU1 enzymatic activity *in vitro*. Purified recombinant NEU1 was incubated with or without purified recombinant CTSA and HS, followed by NEU1 enzymatic activity measurement at pH 4.5. Data represent moles of substrate per moles of NEU1 per second and show means (±SD) of triplicate experiments. Graph in the insert shows expanded view of the results obtained with the apo form of NEU1. (**K)** Docked pose of a HS tetramer in the active site of NEU1 (8DU5). Docking was performed with Autodock. The arginine triad of NEU1 is able to engage sulfate and carboxylate groups of the ligand.

NEU1, together with three other lysosomal enzymes, Cathepsin A (CTSA), β-galactosidase (GLB1) and N-acetylgalactosamine 6-sulfatase (GALNS), forms the lysosomal multienzyme complex (LMC) [53,54]. CTSA protects both GLB1 and NEU1 from lysosomeal degradation, and activates NEU1 (reviewed in ref. [55]), while LMC dissociation evoked by CTSA mutations leads to degradation of both NEU1 and GLB1 [56] and causes, in humans, the neurological LSD galactosialidosis.

Consistent with the assumption that the observed NEU1 deficiency in neurological MPS is caused by LMC dissociation, GLB1 and GALNS activities were reduced in all analysed samples of MPS patients to 25%-50% of the corresponding controls (Figure 4C, D). Unlike the human enzymes, mouse GLB1 and GALNS appear to be stable and active in the absence of CTSA[57,58]. Indeed, in brain tissues of 4-month-old *Hgsnat^P304L^* and *Hgsnat-Geo* mice, only a small, although statistically significant, reduction in GALNS activity was observed (Figure 4E). This finding is reminiscent of the situation in *CathA^S190A-neo^* mice, the *Ctsa* hypomorph mouse model of galactosialidosis [58], where NEU1 activity in the brain tissues was reduced to below 10% of normal, whereas GALNS and GLB1 activities were decreased by only ∼30% and ∼20%, respectively (Figure 4F).

In the brains of human MPS patients and MPS IIIC mice, carboxypeptidase activity measured with the specific CTSA substrate CBZ-Phe-Leu was not reduced (Figure 4G and 4H), ruling out that absence or deficiency of the CTSA protein caused the LMC disruption and secondary NEU1 deficiency in these cases. The LMC integrity was further assessed by size-exclusion chromatography analysis of protein brain extracts. The gel-filtration profiles of NEU1 and GLB1 activities were compared for the WT, *Hgsnat^P304L^*, and galactosialidosis *CathA^S190A-Neo^*mice. For the WT brain, both the intact ∼1,200 kDa LMC and 240 kDa GLB1 tetramers [54] were detected by peaks of GLB1 enzymatic activity eluting from the column at retention times corresponding to ∼1,200 kDa and 240 kDa, respectively. The majority of NEU1 activity was associated with the ∼1,200 kDa LMC peak (Figure 4I, left panel). A major peak for the GLB1 tetramer, with no or small GLB1 and NEU1 activity peaks, corresponding to the elution volume of LMC, was detected in fractions from *CathA^S190A-Neo^*and *Hgsnat^P304L^* mice, respectively, indicating a dissociation of the LMC (Figure 4I, middle and right panels).

*HS inhibits NEU1 activity and causes precipitation of CTSA, GBL1 and NEU1 proteins in vitro.* We further tested if HS would interfere with the activity of the recombinant purified NEU1 or NEU1-CTSA complex in vitro. The proteins were produced in the Sf9 insect cells and purified as previously described [33,34]. Both hexahistidine-tagged and untagged CTSA and NEU1 proteins were tested to exclude the possibility of an unspecific interaction between HS and the positively charged tag. Our results demonstrated that high HS concentrations (0.3 mg/ml and more) reduced the activity of NEU1 in the complex with CTSA (Figure 4J) approximately 100 fold i.e. to the levels of NEU1 not bound to CTSA. A similar effect was also observed for the unbound apo form of NEU1 (Figure 4J insert). Moreover, incubation of CTSA, GLB1 and NEU1 at equimolar amounts with 1 mg/mL of purified HS at pH 4.5/100 mM NaCl, mimicking the conditions in the lysosomal lumen, caused precipitation of all three proteins, either isolated or in combination, suggesting that HS directly binds to each of them and interferes with the correct structural organization of their complex (Figure S4). Notably, hyaluronan, an anionic but not sulphated glycosaminoglycan, did not inhibit NEU1 or caused its precipitation even at the highest studied concentration of 1 mg/ml (not shown).

Based on these findings we conducted molecular docking of a HS tetramer to the NEU1 active site (AutoDock Vina, version 1.2.5) [36], which identified poses where the aminosulphate and carboxylate groups of HS could engage with the basic residues of the Arg triad in NEU1 and also gain additional contacts in the active site (Figure 4K). The pKa of the iduronic acid carboxylate is known to be elevated relative to that of sialic acid (pKa = 3-5 vs. 2.6)[59]. Docking at the CTSA/GLB1 interface did not identify a distinct binding site due to a shallower binding pocket. Further experimental support is required to confirm the specific interactions of HS with these proteins.

### Sialylation of N-linked glycans of brain glycoproteins but not polysialylation of NCAM is increased in MPS patients and mice

NEU1 normally cleaves sialic acid residues from glycans of glycoproteins either during lysosomal breakdown or at the cell surface (reviewed in ref. [60]). Therefore, to assess the effect of NEU1 deficiency on glycoprotein sialylation in MPS brains, N-linked brain glycans were analysed by MALDI-TOF mass spectrometry. Compared to controls, multiple species of sialoglycans were elevated in both *Hgsnat^P304L^*and *Hgsnat-Geo* mice, especially multiple isomeric species containing 2 or more sialic acid residues (Figure 5A). Similarly, highly sialylated glycans, particularly di-, tri- and tetra-sialoglycans, were increased in the samples from MPS IIIA, IIIC and IIID patients (Figure 5B). In particular, a strong increase in a tetra-sialylated species at *m/z* 4761.3 (circled in red in Figure 5B) was observed in MPS IIIC. MS/MS analysis of these species revealed fragments consistent with at least four distinct isomeric structures differing in the position of terminal sialic acid and fucose units (data not sown). In MPS I and II, a general increase in sialylated species was detected together with a widespread increase in complex fucosylated species compared to oligomannose structures (Figure 5C and 5D). In MPS I, MS/MS analysis of the increased sialo-glycans at *m/z* 4213.1 and 4574.3 revealed an enhanced content of glyco-isomers bearing antennary sialyl-Lewis epitopes (circled in red in Figure 5C) uncommon in the human brain. In MPS II, two minor unfucosylated glycoforms, at *m/z* 2431.2 and 2792.3 (circled in red in Figure 5D) with one or two antennas each bearing a sialic acid residue, were markedly increased. Taken together, these data indicate that the secondary deficiency of NEU1 leads to oversialylation of brain glycans in all studied cases of neurological MPS.

**Figure 5.**
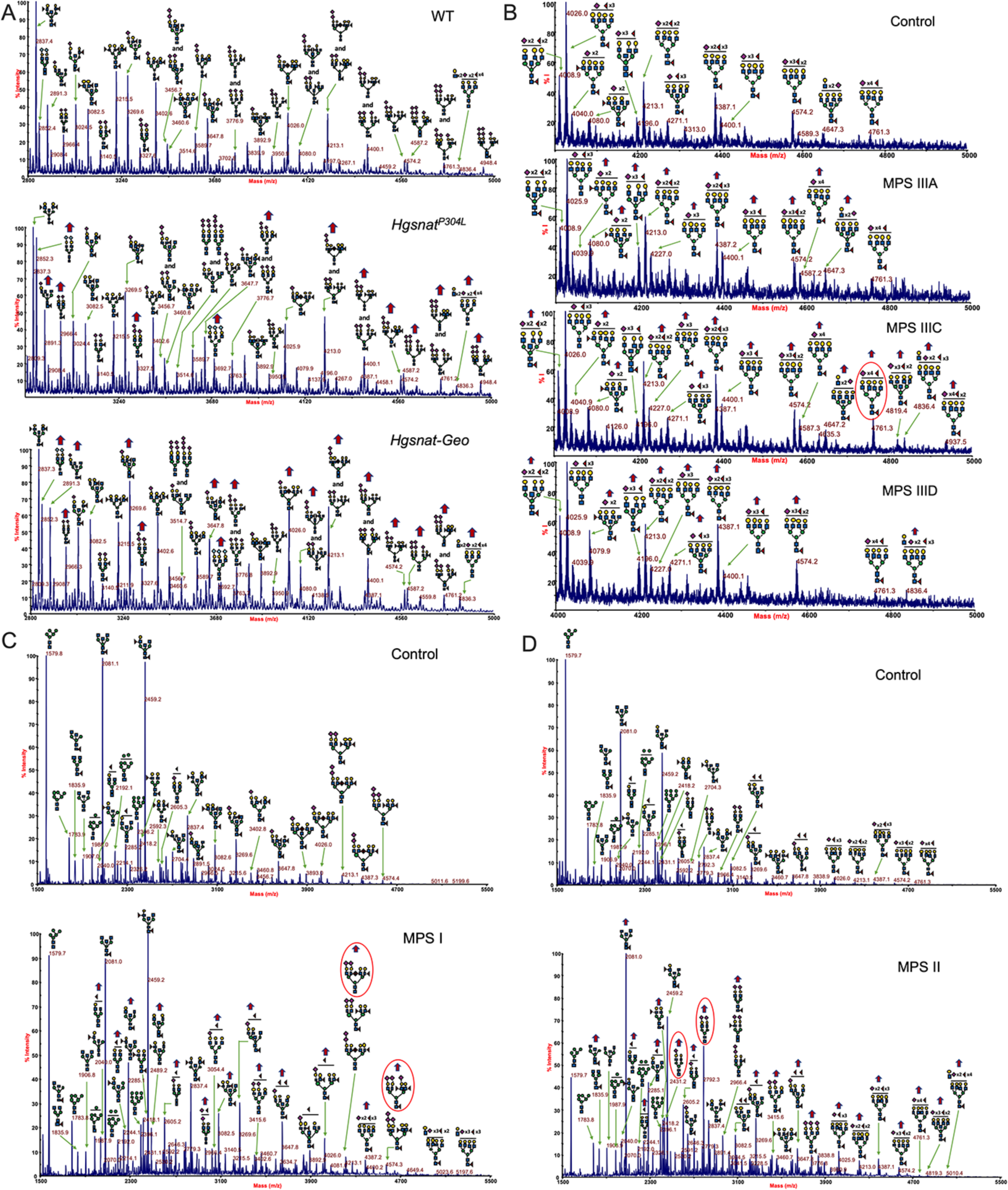
MALDI-TOF mass spectrometry analysis shows increased sialylation of brain proteins in MPS mouse models and human MPS patients. **(A)** Sialylation of N-linked glycans of brain glycoproteins from 10-month-old control, *Hgsnat^P304L^* and *Hgsnat-Geo* mice was analyzed by MALDI-TOF MS. Red arrows mark glycan species containing one or more sialic acid residues with increased intensity for MPS IIIC animals. Each spectrum shows a mass range between m/z 2800 and 5500. **(B)** Sialylation of N-linked glycans from brain glycoproteins of MPS IIIA, IIIC and IIID patients, as well as age and sex-matched controls, was analyzed as in **A**. Mass spectra in the mass range between *m/z* 4000 and 5000 m/z show an increased intensity for sialylated glycan species for MPS III patients (red arrows). **(C)** Sialylation of N-linked glycans from brain glycoproteins of MPS I patient, as well as age and sex-matched control, was analyzed as in **A**. Mass spectra in the range between *m/z* 1500 to 5000 show increased levels of sialylated glycan species for MPS I patients compared to controls (red arrows). Spectra also show a widespread increase in fucosylation together with major increase of glycans bearing sialyl-lewis epitopes (red circles), typically not present in human brain N-glycans. **(D)** Sialylation of N-linked glycans from brain glycoproteins of neurological MPS II patient, as well as age and sex-matched control, was analyzed as in **A**. Mass spectra in the range between *m/z* 1500 to 5000 show increase in sialylated and fucosylated species (red arrows) in MPS II patients. This includes two biantennary unfucosylated N-sialoglycans (circled in red), usually present at a very low level in the human brain N-glycome. GlcNAc, blue square; Man, green circle; Gal, yellow circle; Neu5Ac, purple diamond; Neu5Gc, light blue diamond; Fuc, red triangle.

In contrast to the hypersialylation of other N-linked glycans, a drastic reduction in the polysialylation of NCAM was detected in samples from the anterior cortex of both *Hgsnat^P304L^* and *Hgsnat-Geo* but not *Neu1^−/−^* mice, when measured by a polySia-NCAM-specific sandwich ELISA (Figure 6A). Analysis of the same set of samples by Western blot with polySia-specific antibodies revealed a reduction of the high molecular weight smear bands, characteristic for the polysialylated NCAM isoforms [42,61]. The NCAM-180 protein band detected with NCAM-specific antibodies was also significantly reduced, while NCAM-140 and NCAM-120 protein bands remained at the WT levels (Figure 6C-F). Together, these findings demonstrate that polySia, unlike mono, di-, and oligo-sialylated N-glycans, are not increased by reduced NEU1 activity.

**Figure 6.**
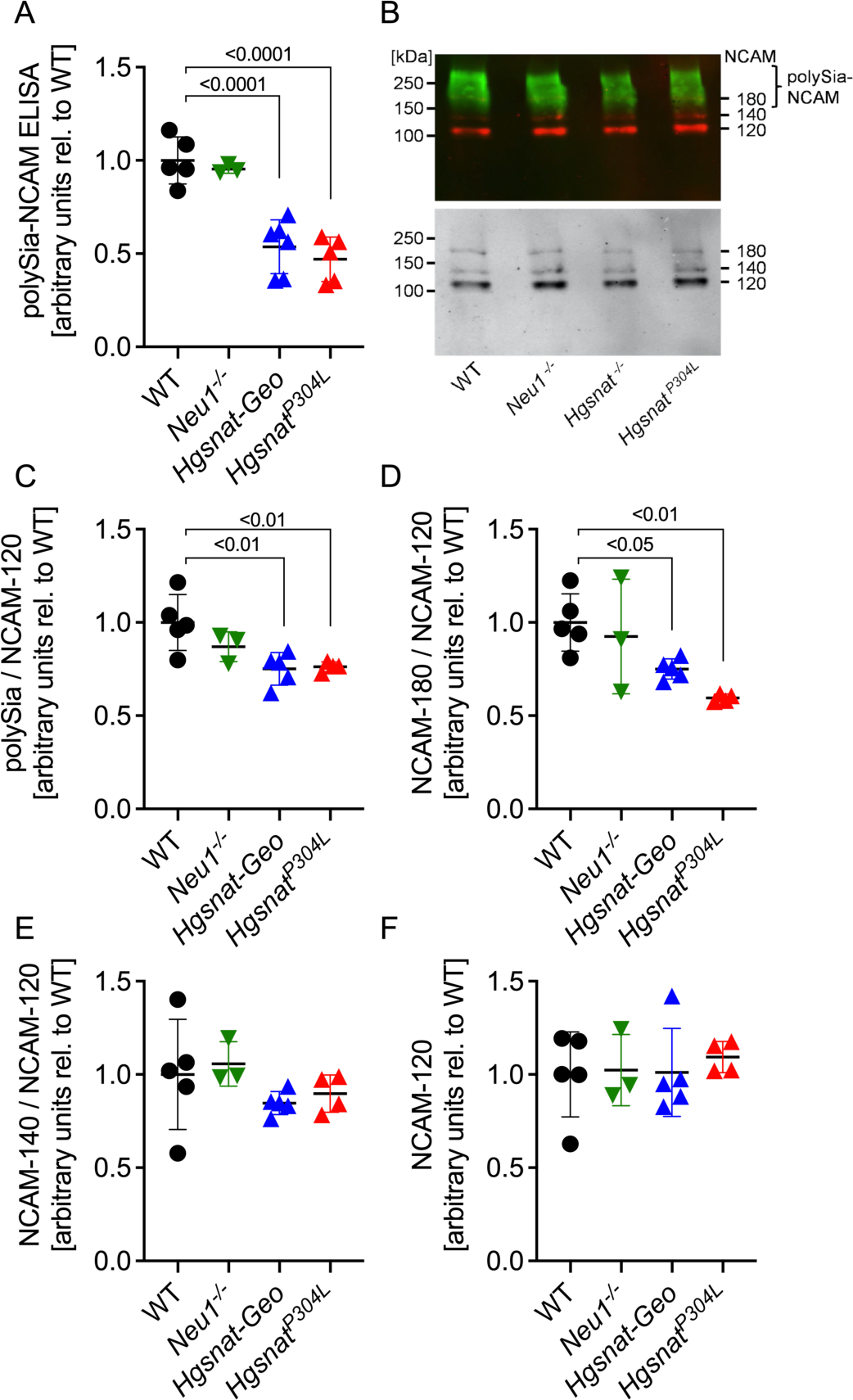
ELISA and Western blot analyses show reduced polySia-NCAM in the cortex of MPS IIIC mouse models. **(A)** PolySia-NCAM levels in lysates of the anterior cortex from 4-month-old WT, *Neu1^−/−^*, *Hgsnat*^−^-*Geo* and *Hgsnat^P304L^* mice were assessed by sandwich ELISA. Values were normalized to the mean values of the WT controls. For each sample from one animal, four technical replicates were measured in independent experiments, and individual values with means (±SD) of 3-6 animals per genotype are plotted. **(B)** Example of Western blot detection with polySia-(800 nm, green, upper panel) and NCAM-specific antibodies (700 nm, red, upper panel) and greyscale image (lower panel). **(C-F)** Densitometric evaluation of polySia-NCAM Western blot signals (**C**) and the bands characteristic for the polySia-free NCAM-140 and NCAM-180 isoforms (**D**, **E**), normalized to the mean values of the WT controls. Values were normalized to the intensities of NCAM-120, the isoform of mature oligodendrocytes, which is not a subject to polysialylation in the mouse brain [61]. (**F**). Individual values with means (±SD) of 3-6 animals per genotype are plotted.

### Transduction of human iPSC-derived cultured cortical MPS IIIA neurons with LV-CTSA-IRES-NEU1-GFP, overexpressing NEU1, partially rescues synaptic defects

To test whether reduction of NEU1 and increased sialylation of brain glycoproteins contribute to synaptic defects in MPS III patients and mouse models [14,62–64], we analyzed peri-axonal puncta positive for post-synaptic protein PSD-95 in juxtaposition with puncta positive for presynaptic protein VGLUT1 in cultured cortical neurons differentiated from induced pluripotent stem cells (iPSCs) derived from a healthy control or from an MPS IIIA patient. NEU1, GALNS, and GLB1 activities as well as densities of juxtapositioned PSD-95- and VGLUT1-positive puncta were drastically reduced in MPS IIIA neurons compared to healthy control (Figure 7A, B). To test if these defects are corrected by restoring deficient levels of NEU1 we transduced MPS IIIA neurons with previously described lentiviral bicistronic vector LV-CTSA-IRES-NEU1-GFP co-expressing human GFP-tagged-NEU1 and CTSA (overexpression of the NEU1 without CTSA results in its aggregation in the cytoplasm [65]). After lentiviral transduction of MPS IIIA neurons with LV-CTSA-IRES-NEU1-GFP, but not with control LV-GFP virus, densities of juxtapositioned PSD-95- and VGLUT1-positive puncta were significantly increased (Figure 7B). Since NEU1 was reported to induce the release of brain-derived neurotrophic factor (BDNF) [66], BDNF levels in the neurons were assessed by immunostaining. BDNF levels in MPS IIIA neurons were below control levels, and not affected by transduction with LV-CTSA-IRES-NEU1-GFP or LV-GFP (Figure 7B, right panel).

**Figure 7.**
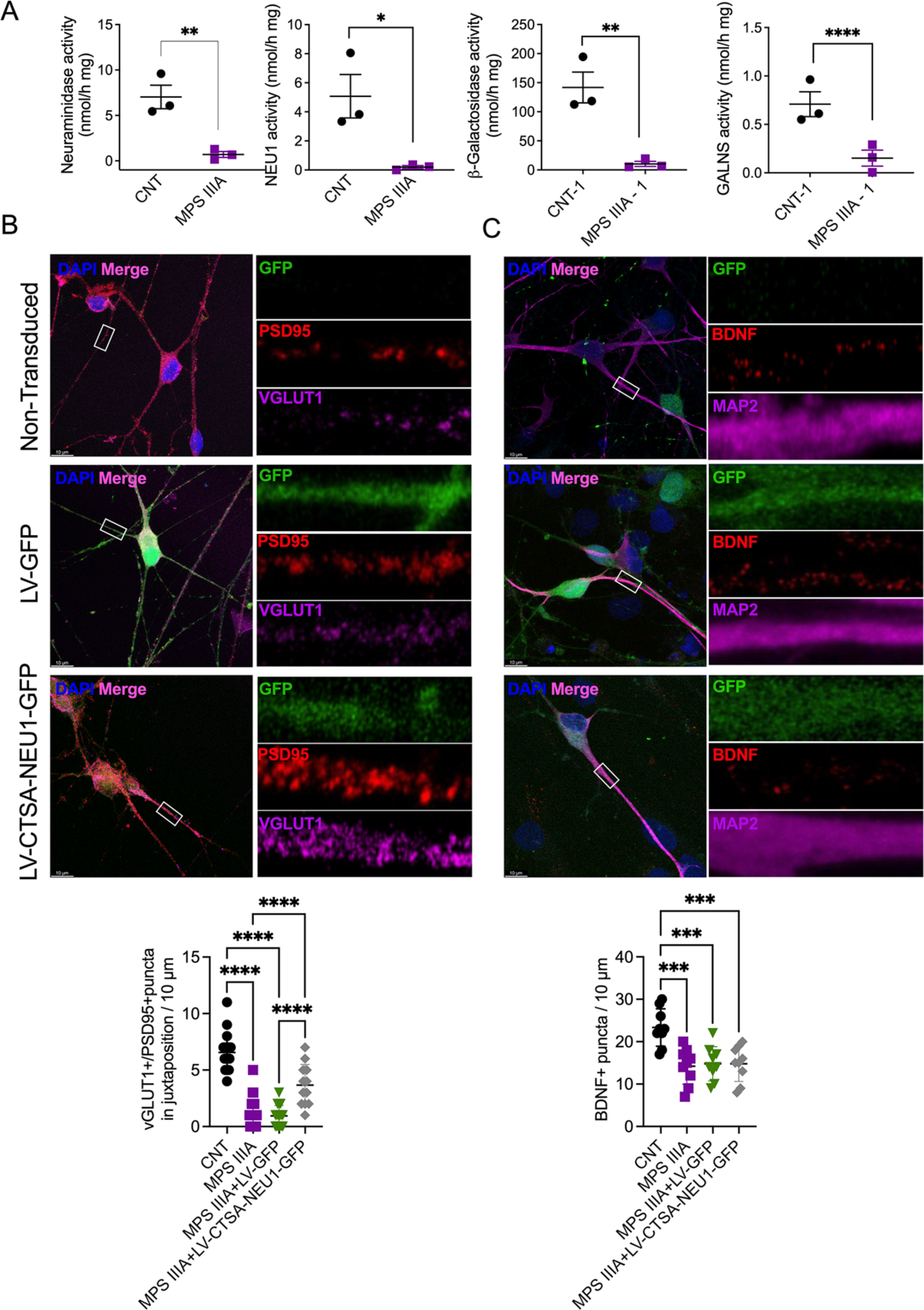
Transduction of human cultured cortical MPS IIIA neurons with LV-CTSA-IRES-NEU1-GFP rescues deficient levels of glutamatergic synapses. **(A)** NEU1, GLB1 and GALNS activities are significantly reduced in cultured iPSC-derived cortical neurons of MPS IIIA patients compared to cells from a healthy control (CNT). The graphs show individual data, means ±SD, n=3, with significance determined by the unpaired t-test. **(B)** MPS IIIA iPSC-derived neurons transduced with LV-CTSA-IRES-NEU1-GFP show higher densities of PSD-95+ puncta in juxtaposition with VGLUT1+ puncta compared to non-transduced or LV-GFP transduced MPS IIIA cells, revealing the increase in the number of functional synapses. **(C)** The amount of BDNF+ puncta (red) is reduced in non-transduced MPS IIIA neurons or those transduced with LV-GFP and LV-CTSA-IRES-NEU1-GFP compared to control. Confocal images were generated by combining 10 Z-stacks, taken at a distance of 0.5 µm using a 63x objective and 2x digital zoom. Scale bars: 10 µm. Puncta were counted manually along the axon at 30 µm increments starting 10 µm from the soma and averaged to 10 µm. Graphs show means ± SD, with significance determined by one-way ANOVA and a Tukey post hoc test.

### Stereotaxic injection of LV-CTSA-IRES-NEU1-GFP in the brain of MPS IIIC mice ameliorates behaviour abnormalities and CNS pathology

To test whether a rescue of NEU1 activity in the brain could also ameliorate known behavioural deficits of MPS IIIC mice [4,14], we performed a bilateral injection of LV-CTSA-IRES-NEU1-GFP in the hippocampi and cortices of P18-P19 *Hgsnat^P304L^* mice. Based on pilot experiments, we determined injection coordinates as 1.5 mm M/L, 2.2 mm A/P, 2.2 mm D/V-, and 1.2-mm D/V to be optimal for targeting the CA1 area of the hippocampus and the layers IV and V of the SS cortex.

Six months after the treatment, we analysed the behaviour of LV-CTSA-IRES-NEU1-GFP injected, sham LV-GFP injected and control WT and *Hgsnat^P304L^* mice using a Novel Object Recognition (NOR) test to assess short-term memory, and Open Field (OF) test to study activity and anxiety levels, as we previously reported [14]. In NOR test with an interval of 1 h between encoding and retrieval, both treated and untreated WT mice showed a positive discrimination index, indicating that they spent more time exploring a novel object in the retrieval phase and compatible with uncompromised short-term recognition memory (Figure 8A). For untreated or sham-treated *Hgsnat^P304L^* mice, the discrimination index was close to zero, and the time spent with the novel object was reduced. However, for *Hgsnat^P304L^*mice injected with LV-CTSA-IRES-NEU1-GFP, both parameters were similar to those of WT mice, suggesting a rescue of the short-term memory deficit (Figure 8A).

**Figure 8.**
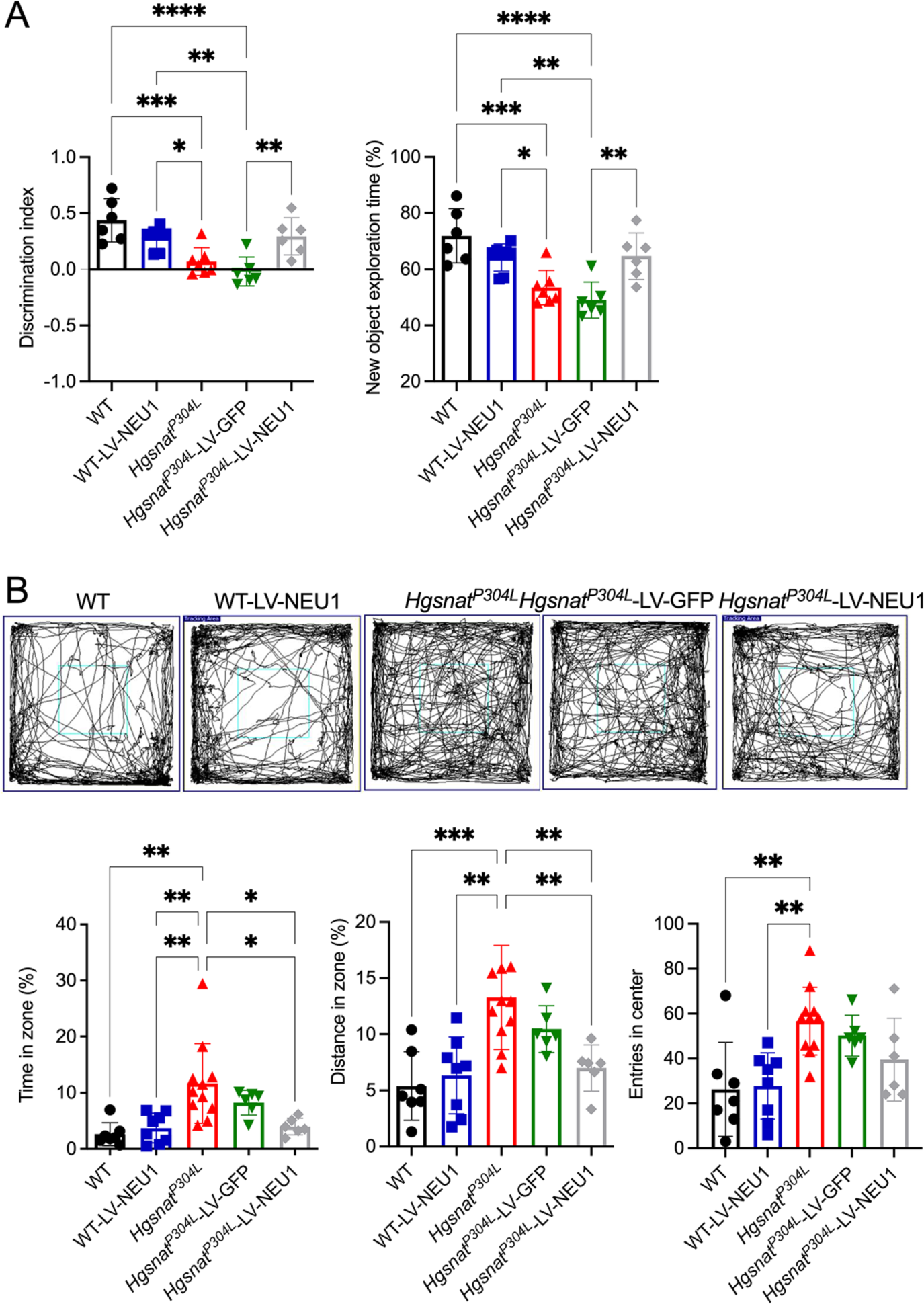
*Hgsnat^P304L^* mice treated with LV-CTSA-IRES-NEU1-GFP show a partial rescue of behaviour abnormalities. **(A)** Improvement of the short-term memory in *Hgsnat^P304L^* mice injected with LV-CTSA-IRES-NEU1-GFP. Discrimination index and the percentage of time spent exploring the novel object were measured using the NOR test in 6-month-old WT mice, WT mice injected with LV-CTSA-IRES-NEU1-GFP, *Hgsnat^P304L^* mice, and *Hgsnat^P304L^* mice injected with LV-CTSA-IRES-NEU1-GFP or LV-GFP. *Hgsnat^P304L^* mice injected with LV-CTSA-IRES-NEU1-GFP show higher discrimination index and time spent with the new object than control or sham treated *Hgsnat^P304L^*mice. **(B)** *Hgsnat^P304L^* mice injected with LV-CTSA-IRES-NEU1-GFP show normalization of anxiety as revealed by the OF test. Graphs show the percentage of time spent in the center of the arena, the percentage of the distance traveled in the center zone and the number of entries to the center of the arena. All graphs show individual data, and means ± SD, n=6-11. Significance was determined by one-way ANOVA and a Tukey post hoc test.

In the OF test, untreated *Hgsnat^P304L^* mice of both sexes spent significantly more time and travelled longer distances in the center zone compared to untreated or treated WT mice (Figure 8B). In contrast, time spent and distance travelled in the center zone for both male and female *Hgsnat^P304L^* mice injected with LV-CTSA-IRES-NEU1-GFP (but not with LV-GFP) was reduced to WT levels consistent with normalization of anxiety (Figure 8B).

Histological analysis of mice following the behavioral tests revealed a widespread expression of both GFP and NEU1-GFP proteins in the neurons of hippocampus, especially the granule cells of the dentate gyrus, the hilar mossy cells, and the CA2/CA3 areas (Figure S5), but not in the neurons of SSC. Such a distribution was not surprising considering that the lentivirus infects mainly dividing or young cells which are found in the dentate gyrus of the hippocampal formation, but not in the SSC (reviewed in ref. [67]).

To test for a rescue of synaptic defects, we analyzed VGLUT1, PSD-95, and Syn1. In agreement with previous findings [14], levels of VGLUT1, PSD-95, and Syn1 puncta in the hippocampi of *Hgsnat^P304L^*and the sham treated *Hgsnat^P304L^* mice were significantly decreased compared to WT mice. Injection of LV-CTSA-IRES-NEU1-GFP did not alter levels of the synaptic proteins in WT mice, but led to a significant increase of VGLUT1 and PSD-95 in *Hgsnat^P304L^* mice, while the levels of Syn1 showed a non-significant trend towards an increase (Figure 9). Consistent with the findings for whole brain extracts (Figure 5A), MALDI-TOF MS and MS/MS analysis indicated increased sialylation of N-linked glycans in the hippocampus of untreated *Hgsnat^P304L^*compared to WT mice, which was normalized by LV-CTSA-IRES-NEU1-GFP injection (Figure S6A). Notably, MS/MS analysis revealed that only 2,3 or 2,6-linked sialic acids were increased in untreated *Hgsnat^P304L^* mice and decreased in LV-CTSA-IRES-NEU1-GFP injected *Hgsnat^P304L^* mice suggesting that NEU1 was active against these types of linkage (Figure S6B). In contrast, glycans bearing 2,8-linked sialic acids were either unaffected or showed a trend opposite to those containing 2,3 or 2,6-linkages. This is consistent with unaltered levels of 2,8-linked polySia in *Neu1^−/−^* mice (Figure 6).

**Figure 9.**
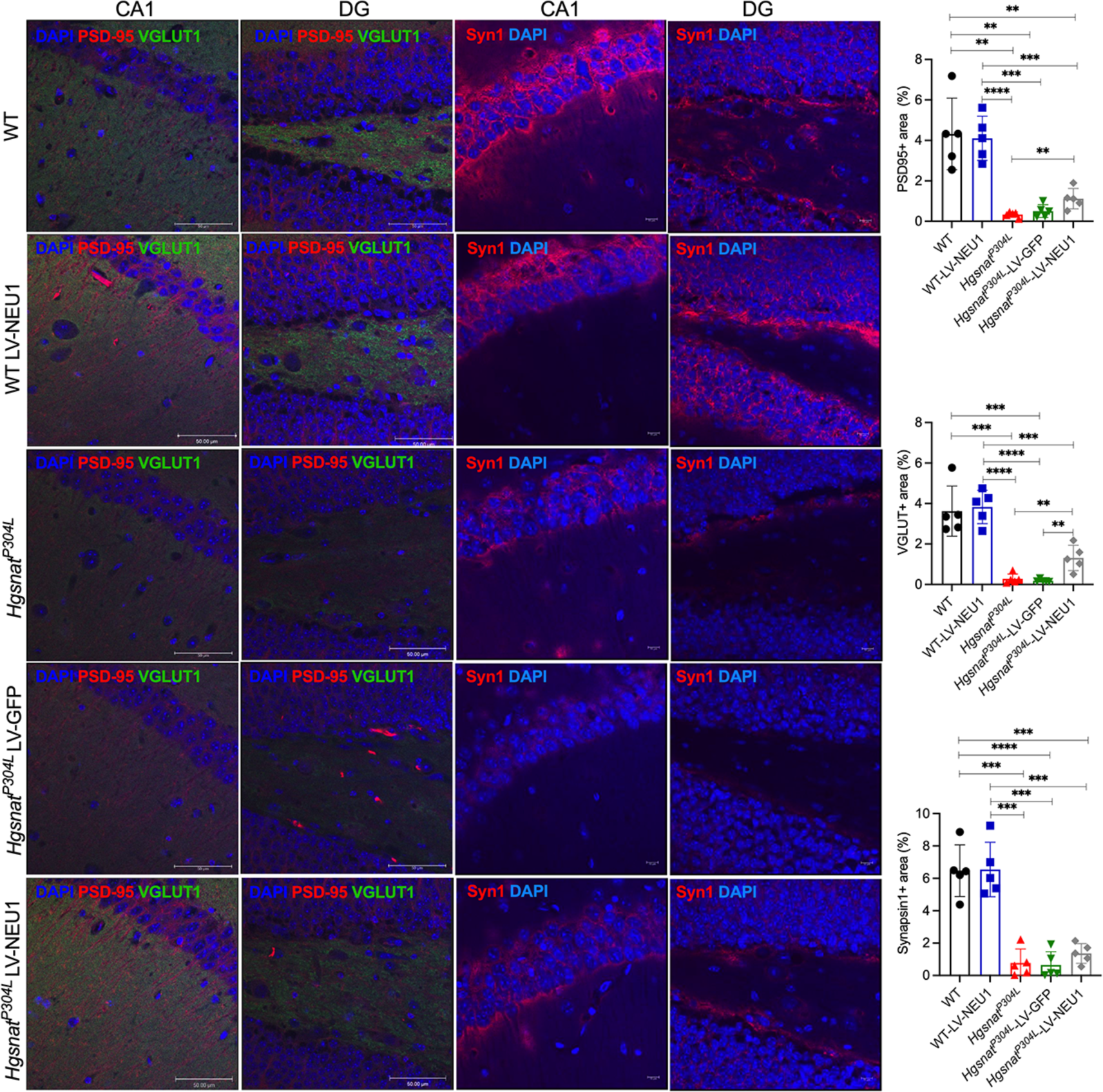
Levels of protein markers of glutamatergic synapse are increased in the brains of *Hgsnat^P304L^* mice injected with LV-CTSA-IRES-NEU1-GFP. Representative confocal images and quantification of fluorescence in four regions of the hippocampus of WT mice, WT mice injected with LV-CTSA-IRES-NEU1-GFP (LV-NEU1), *Hgsnat^P304L^* mice, *Hgsnat^P304L^*mice injected with LV-GFP, and *Hgsnat^P304L^* mice injected with LV-CTSA-IRES-NEU1-GFP. The sections were labeled for PSD-95 (red) and VGLUT1 (far red/pseudo green) or for Syn1 (red) as indicated. The nuclei were counterstained with DAPI (blue). Quantification of images demonstrate that LV-CTSA-IRES-NEU1-GFP treatment increases the levels of both PSD-95 and VGLUT1 in *Hgsnat^P304L^* mice, while Syn1 shows a nonsignificant trend for an increase. Scale bars: 50 and 10 µm. Data on the graphs show individual results, and means (± SD), n=5. Significance was determined by unpaired t-test with multiple correction.

Amyloid deposits are among the hallmarks of MPS III [4,5,7,68] and are associated with increased amyloid beta secretion in *Neu1* KO mice [69]. Consistent with these findings, β-amyloid protein levels were increased in *Hgsnat^P304L^* mice, but unaffected by LV-CTSA-IRES-NEU1-GFP injection (Figure S7). We also found no changes in the levels of GM2 ganglioside, one of the main secondary storage materials in the brains of MPS III patients and mice (Figure S8).

## Discussion

The present study reveals a previously unknown pathophysiological pathway in neurological MPS diseases caused by accumulation of HS, manifesting with a secondary deficiency of NEU1 and leading to synaptic defects. While studying the secondary changes in the activities of lysosomal enzymes in the brain tissues of *Hgsnat^P304L^* and *Hgsnat-Geo* MPS IIIC mouse strains, we serendipitously discovered a ∼50% reduction in total neuraminidase activity in both models. A reduction of neuraminidase activity was also reported for the brain tissues of MPS IIID goats and MPS IIIB mice but remained unexplained [13,70]. An even more drastic decrease was observed in the activity of the lysosomal NEU1 isoform, measured in the presence of the specific inhibitor of NEU3 and NEU4, C9-4BPT-DANA [28]. Since increased lysosomal biogenesis, linked to activation of TFEB-driven expression of lysosomal genes [4,9,12,13], occurs in multiple mouse models of neurological MPS, including those of MPS IIIC [4,9,12,14,71–75], NEU1 activity showed a trend opposite to that of other lysosomal enzymes and proteins.

We further found a correlation between the levels of HS storage and the reductions of NEU1 activity and protein levels in different brain regions of MPS IIIC mice. In particular, NEU1 was deficient in the hippocampus and SSC, the areas with high storage of HS, but not in the olfactory bulb, where HS was not accumulated. Likewise, the pronounced ∼80% decrease of NEU1 activity in liver and lungs coincided with high levels of HS. In contrast, the kidney, an organ with high NEU1 levels but low HS storage [12], showed no NEU1 deficiency.

The secondary NEU1 deficiency occurred in the brains of all analysed mouse models of neurological MPS (MPS I, II, IIIA, IIIB, IIIC), but not in the models of other LSD (Metachromatic leukodystrophy, Mucolipidosis IV, Tay-Sachs or Niemann-Pick type C1). This suggests that NEU1 deficiency is shared only by the lysosomal disorders that store HS. MPS IVA mice, which accumulate mainly keratan sulphate, seem to defy this trend, because they showed a slight but statistically significant reduction in the brain NEU1 activity. However, some MPS IVA patients also display increased tissue levels of HS [76]. Thus, a respective HS storage may occur in the brain of MPS IVA mice and contribute to the partial NEU1 deficiency. Most importantly, our data show that NEU1 deficiency manifests in the brains of human neurological MPS patients, making this finding relevant to our understanding of human pathology.

When BMDM, known to internalize exogenous HS into the lysosomes [77], were treated with urinary HS oligomers resembling those accumulated in the brain [78]. NEU1 activity was reduced proportionally to the HS levels in the culture medium. Reciprocally, AAV-mediated genetic correction of HGSNAT deficiency and HS storage in the brain of *Hgsnat-Geo* mice restored NEU1 activity, proving that NEU1 deficiency was caused by the primary storage of HS and establishing a causal relationship between HS storage and NEU1 deficiency.

The amount of *Neu1* mRNA in the brain tissues of *Hgsnat^P304L^* and *Hgsnat-Geo* mice was similar to WT, while the amount of NEU1 protein in neurons was reduced, suggesting that the mechanism of the deficiency was post-translational, possibly associated with reduced NEU1 stability. A similar phenomenon occurs in the cells of galactosialidosis patients with a genetic deficiency of CTSA leading to disruption of the LMC [54,79]. Notably, the activities of two other LMC components, GLB1 and GALNS, stabilized by their association with CTSA [54] were reduced by at least 50% in brain tissues of MPS patients and cultured iPSC-derived MPS IIIA neurons, while in *Hgsnat^P304L^* mice brain extracts GLB1 was present in the form of homotetramers instead of LMC. Together, these results implicate HS in the dissociation of the complex leading to secondary NEU1, GLB1 and GALNS deficiencies in the brains of neurological MPS patients. In contrast, in MPS IIIC mouse brains, NEU1 was deficient, but GLB1 was increased and GALNS only slightly reduced like in the tissues of galactosialidosis *CathA^S190A-neo^* mice, consistent with high stability of mouse GLB1 and GALNS enzymes in the lysosome [58]. These in vivo results were partially corroborated by in vitro analysis demonstrating that at high HS concentrations caused precipitation of human purified recombinant NEU1, GLB1 and CTSA and the complete loss of NEU1 activity. Computational prediction of specific interactions between HS and proteins can be challenging [80–82]. Based on the data presented here, we hypothesize that HS interacts either with NEU1 or the CTSA/GLB1 protein-protein interface disrupting the correct organization of the LMC and making NEU1 and GLB1 enzymes more susceptible to lysosomal degradation. However, further studies are necessary to identify the underlying structural mechanism and clarify whether storage of HS also causes deficiency of other lysosomal enzymes or proteins.

The level of protein sialylation in the brain is not static, but changes dramatically during development and in response to physiological conditions. Moreover, neuraminidases, including NEU1, were shown to be, at least partially, responsible for these changes (reviewed in ref. [15]). In particular, intracerebral injections of the pan-neuraminidase inhibitor DANA resulted in a significant decrease in LTP, an increase in short-term depression, and an alteration of synaptic plasticity in rodents [83]. Similarly, DANA inhibited long-term potentiation at mossy fibre-CA3 synapses of rats and increased their escape latency in the Morris water maze test [84]. The same group reported that NEU activity was increased after neuronal excitation, and that the removal of sialic acid from the neuronal surface was critical for hippocampal memory formation in a contextual fear-conditioning paradigm [85]. In this study, MS analysis of N-linked glycans revealed that NEU1 deficiency in *Hgsnat^P304L^* and *Hgsnat-Geo* mice, as well as in MPS I, II, and III human patients, was associated with an overall increase in the sialylation of brain glycoproteins which could potentially modify hippocampal synaptic transmission and plasticity.

Consistently, *Hgsnat^P304L^* mice injected with the LV-CTSA-IRES-NEU1-GFP showed higher levels of both pre- and postsynaptic glutamatergic protein markers, VGLUT1 and PSD-95, respectively, in the hippocampus, consistent with amelioration of glutamatergic synaptic defects. The number of Syn1+ puncta also showed a trend for increase in the cells overexpressing NEU1. Similar results were observed in human iPSC-derived cultured MPS IIIA neurons transduced with the LV-CTSA-IRES-NEU1-GFP. Consistent with an increase in the levels of glutamatergic synaptic proteins, we also observed an improvement in short-term memory and partial rescue of reduced anxiety in *Hgsnat^P304L^* mice treated with LV-CTSA-IRES-NEU1-GFP. This partial improvement of behavior could be directly linked to the increased NEU1 activity and amelioration in glutamatergic neurotransmission in the hippocampus, the brain area responsible for short-term memory and modulation of anxiety.

Deficiency of NEU1 in the brain has been previously implicated in amyloidogenesis [69]. *Neu1* KO mice spontaneously developed amyloid plaques presumably due to increased release of hypersialylated amyloid precursor protein. In contrast, plaques were markedly decreased in a mouse model of Alzheimer’s disease treated with AAV9 overexpressing NEU1 [69]. Since amyloidogenesis also occurs in MPS III brain, it could be related to NEU1 deficiency. Our experiments, however, could neither confirm nor reject the link between the NEU1 deficiency and amyloidogenesis in the MPS III. In MPS IIIC mice, accumulation of misfolded amyloid protein occurs mainly in the level 4 and 5 pyramidal neurons of the cortex [61], where we could not observe any expression of the NEU1-GFP after the injection of the LV. Thus, although β-amyloid staining in the neurons of untreated *Hgsnat^P304L^* mice was similar to those treated with LV-CTSA-IRES-NEU1-GFP, we cannot exclude that the absence of the effect was due to the lack of NEU1 expression. In contrast, the hippocampal neurons that showed high transduction with LV-CTSA-IRES-NEU1-GFP do not, in general, reveal any APP accumulation in MPS IIIC. Therefore, further experiments, involving targeted transduction of cortical neurons with the NEU1-expressing virus, should be performed to test the association of NEU1 deficiency with amyloidogenesis. Notably, we also could not detect any reduction in the secondary storage of GM2 ganglioside in the cells overexpressing NEU1, which suggested that the NEU1 deficiency doesn’t contribute to the impairment of ganglioside catabolism and/or autophagy in MPS IIIC neurons.

It has been also proposed that NEU1 plays a role in degradation of polySia attached to NCAM and perhaps other brain proteins. PolySia have been implicated in crucial biological processes in the brain, such as neuronal migration, adhesion and differentiation [86], while altered polysialylation has been associated with severe neurological disorders, such as schizophrenia or Alzheimer’s disease [87]. Specifically, NEU1-catalysed removal of polySia was proposed to participate in lamination of hippocampal granule cells during neuronal migration and in response to stressful events [66,88]. In addition, NEU1 action resulted in secretion of pre-retained BDNF and other neurotrophic molecules, essential to cope with stress conditions, from the polySia chains [66,89]. Our, current data, however, did not reveal an increase in polysialylation of NCAM, the major polySia-modified protein in the brain. In contrast, in the brain of the two MPS IIIC mouse models with secondary Neu1 deficiency, we observed a significant reduction in polySia-NCAM and of the NCAM-180 isoform, both implicated in synapse formation and plasticity [90,91]. Notably, polySia-NCAM was not significantly altered in the *Neu1* KO mice. The alterations in the MPS IIIC mouse models, therefore, are, most likely, not caused by the secondary NEU1 deficiency, but related to neurodegeneration and impaired synaptic transmission observed in these mice [4,14].

To conclude, the current study demonstrates that the secondary deficiency of NEU1 due to the HS-mediated dissociation of the LMC leading to oversialylation of glycoproteins in the brain tissues is a feature common for all neurological MPS diseases. Furthermore, our data show that NEU1 deficiency contributes to the CNS pathology in MPS diseases through defects in synaptic plasticity, yielding novel insights into MPS pathophysiology and suggesting new routes for therapeutic intervention.

## Acknowledgements

We thank Drs Jeffrey A. Medin and Monty McKillop for the help in production of LV-NEU1-GFP and LV-GFP, Drs Volkmar Gieselmann, Shunji Tomatsu and Steven U. Walkley for the precious gift of brain tissues of MLD, MPS IVA, ML IV, and NPC1 mouse models. We also thank Dr. Elke Küster-Schöck and the Plateforme d’Imagerie Microscopique (PIM – CHU Sainte Justine) for the help with confocal microscopy, Annie Nguyen for performing X-Gal staining of brain tissues and Dr. Mila Ashmarina for critically reading the manuscript and helpful advice. Molecular graphics and analyses performed with UCSF ChimeraX, developed by the Resource for Biocomputing, Visualization, and Informatics at the University of California, San Francisco, with support from National Institutes of Health R01-GM129325 and the Office of Cyber Infrastructure and Computational Biology, National Institute of Allergy and Infectious Diseases.

## Authors’ contributions

TM.X., R.H-R., T.M., P.D., X.P., T.G., R.H., L.S., A.P., I.R. and H.T. conducted experiments and acquired data; TM.X., R.H-R., T.M., P.D., L.S., A.P., D.G., I.R., H.T., H.H., C.W.C., T.L. and A.V.P. analysed data; B.B., T.M.D., T.L., J.A., B.A., C.W.C. provided essential resources; A.V.P., D.G., L.S., H.H., TM.X., R.H-R. designed the experiments and wrote the manuscript (first draft); B.B., T.M.D., T.L., J.A., B.A., C.W.C. and A.V.P. edited the manuscript. All authors read and approved the final version of the manuscript.

## Abbreviations

BMDM: bone marrow derived macrophage
CNS: central nervous system
CPC: cetylpyridinium chloride
CTSA: cathepsin A
GAG: glycosaminoglycan
GALNS: glucosamine-6-suphate sulphatase
GLB1: lysosomal β-galactosidase
HGSNAT: acetyl-CoA:alpha-glucosaminide N-acetyltransferase
HS: heparan sulphate
iPSC: induced pluripotent stem cell
KO: knockout
LMC: lysosomal multienzyme complex
LSD: lysosomal storage disease
MALDI-TOF: matrix-assisted laser desorption/ionization – time of flight
MLD: metachromatic leukodystrophy
ML IV: mucolipidosis IV
MPS: mucopolysaccharidosis
NEU1: neuraminidase 1
NPC: neural progenitor cells
NPC1: Niemann-Pick type C1
PolySia: polysialic acid
PolySia-NCAM: polysialylated neural cell adhesion molecule
WT: wild type

## References

1. Neufeld, E.F.; Muenzer, J. The Mucopolysaccharidoses. In The Online Metabolic and Molecular Bases of Inherited Disease, Beaudet, A.L., Vogelstein, B., Kinzler, K.W., Antonarakis, S.E., Ballabio, A., Gibson, K.M., Mitchell, G., Eds. The McGraw-Hill Companies, Inc.: New York, NY, 2014.

2. Heon-Roberts, R.; Nguyen, A.L.A.; Pshezhetsky, A.V. Molecular Bases of Neurodegeneration and Cognitive Decline, the Major Burden of Sanfilippo Disease. J Clin Med 2020, 9, doi:10.3390/jcm9020344.

3. Hamano, K.; Hayashi, M.; Shioda, K.; Fukatsu, R.; Mizutani, S. Mechanisms of neurodegeneration in mucopolysaccharidoses II and IIIB: analysis of human brain tissue. Acta Neuropathol 2008, 115, 547–559, doi:10.1007/s00401-007-0325-3.

4. Martins, C.; Hulkova, H.; Dridi, L.; Dormoy-Raclet, V.; Grigoryeva, L.; Choi, Y.; Langford-Smith, A.; Wilkinson, F.L.; Ohmi, K.; DiCristo, G., et al. Neuroinflammation, mitochondrial defects and neurodegeneration in mucopolysaccharidosis III type C mouse model. Brain 2015, 138, 336–355, doi:10.1093/brain/awu355.

5. Ohmi, K.; Kudo, L.C.; Ryazantsev, S.; Zhao, H.Z.; Karsten, S.L.; Neufeld, E.F. Sanfilippo syndrome type B, a lysosomal storage disease, is also a tauopathy. Proceedings of the National Academy of Sciences of the United States of America 2009, 106, 8332–8337, doi:10.1073/pnas.0903223106.

6. Settembre, C.; Fraldi, A.; Jahreiss, L.; Spampanato, C.; Venturi, C.; Medina, D.; de Pablo, R.; Tacchetti, C.; Rubinsztein, D.C.; Ballabio, A. A block of autophagy in lysosomal storage disorders. Hum Mol Genet 2008, 17, 119–129, doi:10.1093/hmg/ddm289.

7. Viana, G.M.; Priestman, D.A.; Platt, F.M.; Khan, S.; Tomatsu, S.; Pshezhetsky, A.V. Brain Pathology in Mucopolysaccharidoses (MPS) Patients with Neurological Forms. J Clin Med 2020, 9, doi:10.3390/jcm9020396.

8. Ohmi, K.; Greenberg, D.S.; Rajavel, K.S.; Ryazantsev, S.; Li, H.H.; Neufeld, E.F. Activated microglia in cortex of mouse models of mucopolysaccharidoses I and IIIB. Proc Natl Acad Sci U S A 2003, 100, 1902–1907, doi:10.1073/pnas.252784899.

9. Viana, G.M.; Gonzalez, E.A.; Alvarez, M.M.P.; Cavalheiro, R.P.; do Nascimento, C.C.; Baldo, G.; D’Almeida, V.; de Lima, M.A.; Pshezhetsky, A.V.; Nader, H.B. Cathepsin B-associated Activation of Amyloidogenic Pathway in Murine Mucopolysaccharidosis Type I Brain Cortex. Int J Mol Sci 2020, 21, doi:10.3390/ijms21041459.

10. Hadfield, M.G.; Ghatak, N.R.; Nakoneczna, I.; Lippman, H.R.; Myer, E.C.; Constantopoulos, G.; Bradley, R.M. Pathologic Findings in Mucopolysaccharidosis Type IIIB (Sanfilippo’s Syndrome B). Archives of Neurology 1980, 37, 645–650, doi:10.1001/archneur.1980.00500590069012.

11. Heuer, G.G.; Passini, M.A.; Jiang, K.; Parente, M.K.; Lee, V.M.; Trojanowski, J.Q.; Wolfe, J.H. Selective neurodegeneration in murine mucopolysaccharidosis VII is progressive and reversible. Ann Neurol 2002, 52, 762–770, doi:10.1002/ana.10373.

12. Bhaumik, M.; Muller, V.J.; Rozaklis, T.; Johnson, L.; Dobrenis, K.; Bhattacharyya, R.; Wurzelmann, S.; Finamore, P.; Hopwood, J.J.; Walkley, S.U., et al. A mouse model for mucopolysaccharidosis type III A (Sanfilippo syndrome). Glycobiology 1999, 9, 1389–1396.

13. Li, H.H.; Yu, W.H.; Rozengurt, N.; Zhao, H.Z.; Lyons, K.M.; Anagnostaras, S.; Fanselow, M.S.; Suzuki, K.; Vanier, M.T.; Neufeld, E.F. Mouse model of Sanfilippo syndrome type B produced by targeted disruption of the gene encoding alpha-N-acetylglucosaminidase. Proceedings of the National Academy of Sciences of the United States of America 1999, 96, 14505–14510.

14. Pan, X.; Taherzadeh, M.; Bose, P.; Heon-Roberts, R.; Nguyen, A.L.A.; Xu, T.; Para, C.; Yamanaka, Y.; Priestman, D.A.; Platt, F.M., et al. Glucosamine amends CNS pathology in mucopolysaccharidosis IIIC mouse expressing misfolded HGSNAT. J Exp Med 2022, 219, doi:10.1084/jem.20211860.

15. Pshezhetsky, A.V.; Ashmarina, M. Keeping it trim: roles of neuraminidases in CNS function. Glycoconj J 2018, 35, 375–386, doi:10.1007/s10719-018-9837-4.

16. Martins, C.; Hůlková, H.; Dridi, L.; Dormoy-Raclet, V.; Grigoryeva, L.; Choi, Y.; Langford-Smith, A.; Wilkinson, F.L.; Ohmi, K.; DiCristo, G., et al. Neuroinflammation, mitochondrial defects and neurodegeneration in mucopolysaccharidosis III type C mouse model. Brain 2015, 138, 336–355, doi:10.1093/brain/awu355.

17. Pan, X.; De Aragao, C.B.P.; Velasco-Martin, J.P.; Priestman, D.A.; Wu, H.Y.; Takahashi, K.; Yamaguchi, K.; Sturiale, L.; Garozzo, D.; Platt, F.M., et al. Neuraminidases 3 and 4 regulate neuronal function by catabolizing brain gangliosides. FASEB journal : official publication of the Federation of American Societies for Experimental Biology 2017, 10.1096/fj.201601299R, doi:10.1096/fj.201601299R.

18. Seyrantepe, V.; Lema, P.; Caqueret, A.; Dridi, L.; Bel Hadj, S.; Carpentier, S.; Boucher, F.; Levade, T.; Carmant, L.; Gravel, R.A., et al. Mice Doubly-Deficient in Lysosomal Hexosaminidase A and Neuraminidase 4 Show Epileptic Crises and Rapid Neuronal Loss. PLOS Genetics 2010, 6, e1001118, doi:10.1371/journal.pgen.1001118.

19. Clarke, L.A.; Russell, C.S.; Pownall, S.; Warrington, C.L.; Borowski, A.; Dimmick, J.E.; Toone, J.; Jirik, F.R. Murine mucopolysaccharidosis type I: targeted disruption of the murine alpha-L-iduronidase gene. Hum Mol Genet 1997, 6, 503–511, doi:10.1093/hmg/6.4.503.

20. Muenzer, J.; Lamsa, J.C.; Garcia, A.; Dacosta, J.; Garcia, J.; Treco, D.A. Enzyme replacement therapy in mucopolysaccharidosis type II (Hunter syndrome): a preliminary report. Acta Paediatr Suppl 2002, 91, 98–99, doi:10.1111/j.1651-2227.2002.tb03115.x.

21. Li, H.H.; Yu, W.H.; Rozengurt, N.; Zhao, H.Z.; Lyons, K.M.; Anagnostaras, S.; Fanselow, M.S.; Suzuki, K.; Vanier, M.T.; Neufeld, E.F. Mouse model of Sanfilippo syndrome type B produced by targeted disruption of the gene encoding alpha-N-acetylglucosaminidase. Proc Natl Acad Sci U S A 1999, 96, 14505–14510.

22. Tomatsu, S.; Orii, K.O.; Vogler, C.; Nakayama, J.; Levy, B.; Grubb, J.H.; Gutierrez, M.A.; Shim, S.; Yamaguchi, S.; Nishioka, T., et al. Mouse model of N-acetylgalactosamine-6-sulfate sulfatase deficiency (Galns−/−) produced by targeted disruption of the gene defective in Morquio A disease. Hum Mol Genet 2003, 12, 3349–3358, doi:10.1093/hmg/ddg366.

23. Hess, B.; Saftig, P.; Hartmann, D.; Coenen, R.; Lullmann-Rauch, R.; Goebel, H.H.; Evers, M.; von Figura, K.; D’Hooge, R.; Nagels, G., et al. Phenotype of arylsulfatase A-deficient mice: relationship to human metachromatic leukodystrophy. Proc Natl Acad Sci U S A 1996, 93, 14821–14826, doi:10.1073/pnas.93.25.14821.

24. Matthes, F.; Andersson, C.; Stein, A.; Eistrup, C.; Fogh, J.; Gieselmann, V.; Wenger, D.A.; Matzner, U. Enzyme replacement therapy of a novel humanized mouse model of globoid cell leukodystrophy. Exp Neurol 2015, 271, 36–45, doi:10.1016/j.expneurol.2015.04.020.

25. Maue, R.A.; Burgess, R.W.; Wang, B.; Wooley, C.M.; Seburn, K.L.; Vanier, M.T.; Rogers, M.A.; Chang, C.C.; Chang, T.Y.; Harris, B.T., et al. A novel mouse model of Niemann-Pick type C disease carrying a D1005G-Npc1 mutation comparable to commonly observed human mutations. Hum Mol Genet 2012, 21, 730–750, doi:10.1093/hmg/ddr505.

26. Seyrantepe, V.; Canuel, M.; Carpentier, S.; Landry, K.; Durand, S.; Liang, F.; Zeng, J.; Caqueret, A.; Gravel, R.A.; Marchesini, S., et al. Mice deficient in Neu4 sialidase exhibit abnormal ganglioside catabolism and lysosomal storage. Hum Mol Genet 2008, 17, 1556–1568, doi:10.1093/hmg/ddn043.

27. Potier, M.; Mameli, L.; Bélisle, M.; Dallaire, L.; Melançon, S.B. Fluorometric assay of neuraminidase with a sodium (4-methylumbelliferyl-α-d-N-acetylneuraminate) substrate. Analytical Biochemistry 1979, 94, 287–296, 10.1016/0003-2697(79)90362-2.

28. Guo, T.; Dätwyler, P.; Demina, E.; Richards, M.R.; Ge, P.; Zou, C.; Zheng, R.; Fougerat, A.; Pshezhetsky, A.V.; Ernst, B., et al. Selective Inhibitors of Human Neuraminidase 3. Journal of Medicinal Chemistry 2018, 61, 1990–2008, doi:10.1021/acs.jmedchem.7b01574.

29. Karpova, E.A.; Voznyi Ya, V.; Keulemans, J.L.; Hoogeveen, A.T.; Winchester, B.; Tsvetkova, I.V.; van Diggelen, O.P. A fluorimetric enzyme assay for the diagnosis of Sanfilippo disease type A (MPS IIIA). J Inherit Metab Dis 1996, 19, 278–285, doi:10.1007/bf01799255.

30. Sturiale, L.; Barone, R.; Garozzo, D. The impact of mass spectrometry in the diagnosis of congenital disorders of glycosylation. J Inherit Metab Dis 2011, 34, 891–899, doi:10.1007/s10545-011-9306-8.

31. Palmigiano, A.; Barone, R.; Sturiale, L.; Sanfilippo, C.; Bua, R.O.; Romeo, D.A.; Messina, A.; Capuana, M.L.; Maci, T.; Le Pira, F., et al. CSF N-glycoproteomics for early diagnosis in Alzheimer’s disease. J Proteomics 2016, 131, 29–37, doi:10.1016/j.jprot.2015.10.006.

32. Heap, R.E.; Marin-Rubio, J.L.; Peltier, J.; Heunis, T.; Dannoura, A.; Moore, A.; Trost, M. Proteomics characterisation of the L929 cell supernatant and its role in BMDM differentiation. Life Sci Alliance 2021, 4, doi:10.26508/lsa.202000957.

33. Gorelik, A.; Illes, K.; Mazhab-Jafari, M.T.; Nagar, B. Structure of the immunoregulatory sialidase NEU1. Sci Adv 2023, 9, eadf8169, doi:10.1126/sciadv.adf8169.

34. Gorelik, A.; Illes, K.; Hasan, S.M.N.; Nagar, B.; Mazhab-Jafari, M.T. Structure of the murine lysosomal multienzyme complex core. Sci Adv 2021, 7, doi:10.1126/sciadv.abf4155.

35. Wander, R.; Kaminski, A.M.; Xu, Y.; Pagadala, V.; Krahn, J.M.; Pham, T.Q.; Liu, J.; Pedersen, L.C. Deciphering the substrate recognition mechanisms of the heparan sulfate 3-O-sulfotransferase-3. RSC Chem Biol 2021, 2, 1239–1248, doi:10.1039/d1cb00079a.

36. Eberhardt, J.; Santos-Martins, D.; Tillack, A.F.; Forli, S. AutoDock Vina 1.2.0: New Docking Methods, Expanded Force Field, and Python Bindings. J Chem Inf Model 2021, 61, 3891–3898, doi:10.1021/acs.jcim.1c00203.

37. Kochnev, Y.; Hellemann, E.; Cassidy, K.C.; Durrant, J.D. Webina: an open-source library and web app that runs AutoDock Vina entirely in the web browser. Bioinformatics 2020, 36, 4513–4515, doi:10.1093/bioinformatics/btaa579.

38. Pettersen, E.F.; Goddard, T.D.; Huang, C.C.; Meng, E.C.; Couch, G.S.; Croll, T.I.; Morris, J.H.; Ferrin, T.E. UCSF ChimeraX: Structure visualization for researchers, educators, and developers. Protein Sci 2021, 30, 70–82, doi:10.1002/pro.3943.

39. Piras, F.; Schiff, M.; Chiapponi, C.; Bossu, P.; Muhlenhoff, M.; Caltagirone, C.; Gerardy-Schahn, R.; Hildebrandt, H.; Spalletta, G. Brain structure, cognition and negative symptoms in schizophrenia are associated with serum levels of polysialic acid-modified NCAM. Transl Psychiatry 2015, 5, e658, doi:10.1038/tp.2015.156.

40. Frosch, M.; Gorgen, I.; Boulnois, G.J.; Timmis, K.N.; Bitter-Suermann, D. NZB mouse system for production of monoclonal antibodies to weak bacterial antigens: isolation of an IgG antibody to the polysaccharide capsules of Escherichia coli K1 and group B meningococci. Proc Natl Acad Sci U S A 1985, 82, 1194–1198, doi:10.1073/pnas.82.4.1194.

41. Schulz, E.C.; Dickmanns, A.; Urlaub, H.; Schmitt, A.; Muhlenhoff, M.; Stummeyer, K.; Schwarzer, D.; Gerardy-Schahn, R.; Ficner, R. Crystal structure of an intramolecular chaperone mediating triple-beta-helix folding. Nat Struct Mol Biol 2010, 17, 210–215, doi:10.1038/nsmb.1746.

42. Tantra, M.; Krocher, T.; Papiol, S.; Winkler, D.; Rockle, I.; Jatho, J.; Burkhardt, H.; Ronnenberg, A.; Gerardy-Schahn, R.; Ehrenreich, H., et al. St8sia2 deficiency plus juvenile cannabis exposure in mice synergistically affect higher cognition in adulthood. Behav Brain Res 2014, 275, 166–175, doi:10.1016/j.bbr.2014.08.062.

43. Kawamura, T.; Suzuki, J.; Wang, Y.V.; Menendez, S.; Morera, L.B.; Raya, A.; Wahl, G.M.; Izpisua Belmonte, J.C. Linking the p53 tumour suppressor pathway to somatic cell reprogramming. Nature 2009, 460, 1140–1144, doi:10.1038/nature08311.

44. Chen, X.; Rocha, C.; Rao, T.; Durcan, T. NeuroEDDU protocols_iPSC culture_interactive protocol. 2019, 10.5281/ZENODO.3733913, doi:10.5281/ZENODO.3733913.

45. Chen, X.; Rocha, C.; Loignon, M.; Peng, H.; Rao, T.; Durcan, T.M. Induction of Dopaminergic or Cortical neuronal progenitors from iPSCs. 2019, 10.5281/ZENODO.3738358, doi:10.5281/ZENODO.3738358.

46. Chen, X.; Lauinger, N.; Rocha, C.; Rao, T.; Durcan, T.M. Generation of dopaminergic or cortical neurons from neuronal progenitors (interactive protocol). 2019, 10.5281/ZENODO.3733914, doi:10.5281/ZENODO.3733914.

47. Mooree, T.; Bose, P.; Wood, J.; Durcan, T.; Pshezhetsky, A. iPSC derived neurons of mucopolysaccharidosis III patients show pronounced synaptic defects. Molecular Genetics and Metabolism 2022, 135, S85, 10.1016/j.ymgme.2021.11.219.

48. Fougerat, A.; Pan, X.; Smutova, V.; Heveker, N.; Cairo, C.W.; Issad, T.; Larrivee, B.; Medin, J.A.; Pshezhetsky, A.V. Neuraminidase 1 activates insulin receptor and reverses insulin resistance in obese mice. Mol Metab 2018, 12, 76–88, doi:10.1016/j.molmet.2018.03.017.

49. Fujitsuka, H.; Sawamoto, K.; Peracha, H.; Mason, R.W.; Mackenzie, W.; Kobayashi, H.; Yamaguchi, S.; Suzuki, Y.; Orii, K.; Orii, T., et al. Biomarkers in patients with mucopolysaccharidosis type II and IV. Mol Genet Metab Rep 2019, 19, 100455, doi:10.1016/j.ymgmr.2019.100455.

50. Rowan, D.J.; Tomatsu, S.; Grubb, J.H.; Montano, A.M.; Sly, W.S. Assessment of bone dysplasia by micro-CT and glycosaminoglycan levels in mouse models for mucopolysaccharidosis type I, IIIA, IVA, and VII. J Inherit Metab Dis 2013, 36, 235–246, doi:10.1007/s10545-012-9522-x.

51. Tordo, J.; O’Leary, C.; Antunes, A.; Palomar, N.; Aldrin-Kirk, P.; Basche, M.; Bennett, A.; D’Souza, Z.; Gleitz, H.; Godwin, A., et al. A novel adeno-associated virus capsid with enhanced neurotropism corrects a lysosomal transmembrane enzyme deficiency. Brain 2018, 10.1093/brain/awy126, doi:10.1093/brain/awy126.

52. Ausseil, J.; Desmaris, N.; Bigou, S.; Attali, R.; Corbineau, S.; Vitry, S.; Parent, M.; Cheillan, D.; Fuller, M.; Maire, I., et al. Early neurodegeneration progresses independently of microglial activation by heparan sulfate in the brain of mucopolysaccharidosis IIIB mice. PLoS One 2008, 3, e2296, doi:10.1371/journal.pone.0002296.

53. Pshezhetsky, A.V.; Potier, M. Direct Affinity Purification and Supramolecular Organization of Human Lysosomal Cathepsin A. Archives of Biochemistry and Biophysics 1994, 313, 64–70, 10.1006/abbi.1994.1359.

54. Pshezhetsky, A.V.; Potier, M. Association of N-acetylgalactosamine-6-sulfate sulfatase with the multienzyme lysosomal complex of beta-galactosidase, cathepsin A, and neuraminidase. Possible implication for intralysosomal catabolism of keratan sulfate. J Biol Chem 1996, 271, 28359–28365.

55. Pshezhetsky, A.V.; Ashmarina, M. Lysosomal multienzyme complex: biochemistry, genetics, and molecular pathophysiology. Prog Nucleic Acid Res Mol Biol 2001, 69, 81–114.

56. Vinogradova, M.V.; Michaud, L.; Mezentsev, A.V.; Lukong, K.E.; El-Alfy, M.; Morales, C.R.; Potier, M.; Pshezhetsky, A.V. Molecular mechanism of lysosomal sialidase deficiency in galactosialidosis involves its rapid degradation. Biochem J 1998, 330 (Pt 2), 641–650, doi:10.1042/bj3300641.

57. Zhou, X.Y.; Morreau, H.; Rottier, R.; Davis, D.; Bonten, E.; Gillemans, N.; Wenger, D.; Grosveld, F.G.; Doherty, P.; Suzuki, K., et al. Mouse model for the lysosomal disorder galactosialidosis and correction of the phenotype with overexpressing erythroid precursor cells. Genes Dev 1995, 9, 2623–2634.

58. Seyrantepe, V.; Hinek, A.; Peng, J.; Fedjaev, M.; Ernest, S.; Kadota, Y.; Canuel, M.; Itoh, K.; Morales, C.R.; Lavoie, J., et al. Enzymatic activity of lysosomal carboxypeptidase (cathepsin) A is required for proper elastic fiber formation and inactivation of endothelin-1. Circulation 2008, 117, 1973–1981, doi:10.1161/CIRCULATIONAHA.107.733212.

59. Wang, H.M.; Loganathan, D.; Linhardt, R.J. Determination of the pKa of glucuronic acid and the carboxy groups of heparin by 13C-nuclear-magnetic-resonance spectroscopy. Biochem J 1991, 278 (Pt 3), 689–695, doi:10.1042/bj2780689.

60. Monti, E.; Miyagi, T. Structure and Function of Mammalian Sialidases. In SialoGlyco Chemistry and Biology I: Biosynthesis, structural diversity and sialoglycopathologies, Gerardy-Schahn, R., Delannoy, P., von Itzstein, M., Eds. Springer Berlin Heidelberg: Berlin, Heidelberg, 2015; 10.1007/128_2012_328 pp. 183–208.

61. Oltmann-Norden, I.; Galuska, S.P.; Hildebrandt, H.; Geyer, R.; Gerardy-Schahn, R.; Geyer, H.; Muhlenhoff, M. Impact of the polysialyltransferases ST8SiaII and ST8SiaIV on polysialic acid synthesis during postnatal mouse brain development. J Biol Chem 2008, 283, 1463–1471, doi:10.1074/jbc.M708463200.

62. Para, C.; Bose, P.; Bruno, L.; Freemantle, E.; Taherzadeh, M.; Pan, X.; Han, C.; McPherson, P.S.; Lacaille, J.C.; Bonneil, E., et al. Early defects in mucopolysaccharidosis type IIIC disrupt excitatory synaptic transmission. JCI Insight 2021, 10.1172/jci.insight.142073, doi:10.1172/jci.insight.142073.

63. Sambri, I.; D’Alessio, R.; Ezhova, Y.; Giuliano, T.; Sorrentino, N.C.; Cacace, V.; De Risi, M.; Cataldi, M.; Annunziato, L.; De Leonibus, E., et al. Lysosomal dysfunction disrupts presynaptic maintenance and restoration of presynaptic function prevents neurodegeneration in lysosomal storage diseases. EMBO Mol Med 2017, 9, 112–132, doi:10.15252/emmm.201606965.

64. Dwyer, C.A.; Scudder, S.L.; Lin, Y.; Dozier, L.E.; Phan, D.; Allen, N.J.; Patrick, G.N.; Esko, J.D. Neurodevelopmental Changes in Excitatory Synaptic Structure and Function in the Cerebral Cortex of Sanfilippo Syndrome IIIA Mice. Sci Rep 2017, 7, 46576, doi:10.1038/srep46576.

65. Pshezhetsky, A.V.; Richard, C.; Michaud, L.; Igdoura, S.; Wang, S.; Elsliger, M.A.; Qu, J.; Leclerc, D.; Gravel, R.; Dallaire, L., et al. Cloning, expression and chromosomal mapping of human lysosomal sialidase and characterization of mutations in sialidosis. Nat Genet 1997, 15, 316–320, doi:10.1038/ng0397-316.

66. Abe, C.; Yi, Y.; Hane, M.; Kitajima, K.; Sato, C. Acute stress-induced change in polysialic acid levels mediated by sialidase in mouse brain. Sci Rep 2019, 9, 9950, doi:10.1038/s41598-019-46240-6.

67. Anacker, C.; Hen, R. Adult hippocampal neurogenesis and cognitive flexibility - linking memory and mood. Nat Rev Neurosci 2017, 18, 335–346, doi:10.1038/nrn.2017.45.

68. Ohmi, K.; Zhao, H.Z.; Neufeld, E.F. Defects in the medial entorhinal cortex and dentate gyrus in the mouse model of Sanfilippo syndrome type B. PLoS One 2011, 6, e27461, doi:10.1371/journal.pone.0027461.

69. Annunziata, I.; Patterson, A.; Helton, D.; Hu, H.; Moshiach, S.; Gomero, E.; Nixon, R.; d’Azzo, A. Lysosomal NEU1 deficiency affects amyloid precursor protein levels and amyloid-beta secretion via deregulated lysosomal exocytosis. Nat Commun 2013, 4, 2734, doi:10.1038/ncomms3734.

70. Jones, M.Z.; Alroy, J.; Boyer, P.J.; Cavanagh, K.T.; Johnson, K.; Gage, D.; Vorro, J.; Render, J.A.; Common, R.S.; Leedle, R.A., et al. Caprine mucopolysaccharidosis-IIID: clinical, biochemical, morphological and immunohistochemical characteristics. J Neuropathol Exp Neurol 1998, 57, 148–157, doi:10.1097/00005072-199802000-00006.

71. Baldo, G.; Lorenzini, D.M.; Santos, D.S.; Mayer, F.Q.; Vitry, S.; Bigou, S.; Heard, J.M.; Matte, U.; Giugliani, R. Shotgun proteomics reveals possible mechanisms for cognitive impairment in Mucopolysaccharidosis I mice. Mol Genet Metab 2015, 114, 138–145, doi:10.1016/j.ymgme.2014.12.301.

72. Cardone, M.; Polito, V.A.; Pepe, S.; Mann, L.; D’Azzo, A.; Auricchio, A.; Ballabio, A.; Cosma, M.P. Correction of Hunter syndrome in the MPSII mouse model by AAV2/8-mediated gene delivery. Human molecular genetics 2006, 15, 1225–1236, doi:10.1093/hmg/ddl038.

73. Crawley, A.C.; Gliddon, B.L.; Auclair, D.; Brodie, S.L.; Hirte, C.; King, B.M.; Fuller, M.; Hemsley, K.M.; Hopwood, J.J. Characterization of a C57BL/6 congenic mouse strain of mucopolysaccharidosis type IIIA. Brain Res 2006, 1104, 1–17, doi:10.1016/j.brainres.2006.05.079.

74. Frisella, W.A.; O’Connor, L.H.; Vogler, C.A.; Roberts, M.; Walkley, S.; Levy, B.; Daly, T.M.; Sands, M.S. Intracranial injection of recombinant adeno-associated virus improves cognitive function in a murine model of mucopolysaccharidosis type VII. Mol Ther 2001, 3, 351–358, doi:10.1006/mthe.2001.0274.

75. Garcia, A.R.; Pan, J.; Lamsa, J.C.; Muenzer, J. The characterization of a murine model of mucopolysaccharidosis II (Hunter syndrome). Journal of inherited metabolic disease 2007, 30, 924–934, doi:10.1007/s10545-007-0641-8.

76. Tomatsu, S.; Gutierrez, M.A.; Ishimaru, T.; Pena, O.M.; Montano, A.M.; Maeda, H.; Velez-Castrillon, S.; Nishioka, T.; Fachel, A.A.; Cooper, A., et al. Heparan sulfate levels in mucopolysaccharidoses and mucolipidoses. Journal of inherited metabolic disease 2005, 28, 743–757, doi:10.1007/s10545-005-0069-y.

77. Thomson, M. Endocytosis, partial degradation and release of heparan sulfate by elicited mouse peritoneal macrophages. Int J Biol Macromol 1994, 16, 245–251, doi:10.1016/0141-8130(94)90029-9.

78. Fuller, M.; Rozaklis, T.; Ramsay, S.L.; Hopwood, J.J.; Meikle, P.J. Disease-specific markers for the mucopolysaccharidoses. Pediatr Res 2004, 56, 733–738, doi:10.1203/01.PDR.0000141987.69757.DD.

79. Vinogradova, M.V.; Michaud, L.; Mezentsev, A.V.; Lukong, K.E.; El-Alfy, M.; Morales, C.R.; Potier, M.; Pshezhetsky, A.V. Molecular mechanism of lysosomal sialidase deficiency in galactosialidosis involves its rapid degradation. Biochem J 1998, 330 (Pt 2), 641–650.

80. Kogut, M.M.; Marcisz, M.; Samsonov, S.A. Modeling glycosaminoglycan-protein complexes. Curr Opin Struct Biol 2022, 73, 102332, doi:10.1016/j.sbi.2022.102332.

81. Almond, A. Multiscale modeling of glycosaminoglycan structure and dynamics: current methods and challenges. Curr Opin Struct Biol 2018, 50, 58–64, doi:10.1016/j.sbi.2017.11.008.

82. Capila, I.; Linhardt, R.J. Heparin-protein interactions. Angew Chem Int Ed Engl 2002, 41, 391–412, doi:10.1002/1521-3773(20020201)41:3<390::aid-anie390>3.0.co;2-b.

83. Savotchenko, A.; Romanov, A.; Isaev, D.; Maximyuk, O.; Sydorenko, V.; Holmes, G.L.; Isaeva, E. Neuraminidase inhibition primes short-term depression and suppresses long-term potentiation of synaptic transmission in the rat hippocampus. Neural Plast 2015, 2015, 908190–908190, doi:10.1155/2015/908190.

84. Minami, A.; Saito, M.; Mamada, S.; Ieno, D.; Hikita, T.; Takahashi, T.; Otsubo, T.; Ikeda, K.; Suzuki, T. Role of Sialidase in Long-Term Potentiation at Mossy Fiber-CA3 Synapses and Hippocampus-Dependent Spatial Memory. PLoS ONE 2016, 11, e0165257–e0165257, doi:10.1371/journal.pone.0165257.

85. Minami, A.; Meguro, Y.; Ishibashi, S.; Ishii, A.; Shiratori, M.; Sai, S.; Horii, Y.; Shimizu, H.; Fukumoto, H.; Shimba, S., et al. Rapid regulation of sialidase activity in response to neural activity and sialic acid removal during memory processing in rat hippocampus. J Biol Chem 2017, 292, 5645–5654, doi:10.1074/jbc.M116.764357.

86. Hildebrandt, H.; Dityatev, A. Polysialic Acid in Brain Development and Synaptic Plasticity. Top Curr Chem 2015, 366, 55–96, doi:10.1007/128_2013_446.

87. Thiesler, H.; Kucukerden, M.; Gretenkort, L.; Rockle, I.; Hildebrandt, H. News and Views on Polysialic Acid: From Tumor Progression and Brain Development to Psychiatric Disorders, Neurodegeneration, Myelin Repair and Immunomodulation. Front Cell Dev Biol 2022, 10, 871757, doi:10.3389/fcell.2022.871757.

88. Sajo, M.; Sugiyama, H.; Yamamoto, H.; Tanii, T.; Matsuki, N.; Ikegaya, Y.; Koyama, R. Neuraminidase-Dependent Degradation of Polysialic Acid Is Required for the Lamination of Newly Generated Neurons. PLoS ONE 2016, 11, e0146398–e0146398, doi:10.1371/journal.pone.0146398.

89. Sumida, M.; Hane, M.; Yabe, U.; Shimoda, Y.; Pearce, O.M.; Kiso, M.; Miyagi, T.; Sawada, M.; Varki, A.; Kitajima, K., et al. Rapid Trimming of Cell Surface Polysialic Acid (PolySia) by Exovesicular Sialidase Triggers Release of Preexisting Surface Neurotrophin. J Biol Chem 2015, 290, 13202–13214, doi:10.1074/jbc.M115.638759.

90. Schuster, T.; Krug, M.; Hassan, H.; Schachner, M. Increase in proportion of hippocampal spine synapses expressing neural cell adhesion molecule NCAM180 following long-term potentiation. J Neurobiol 1998, 37, 359–372, doi:10.1002/(sici)1097-4695(19981115)37:3<359::aid-neu2>3.0.co;2-4.

91. Dityatev, A.; Dityateva, G.; Sytnyk, V.; Delling, M.; Toni, N.; Nikonenko, I.; Muller, D.; Schachner, M. Polysialylated neural cell adhesion molecule promotes remodeling and formation of hippocampal synapses. J Neurosci 2004, 24, 9372–9382, doi:10.1523/JNEUROSCI.1702-04.2004.

