## Supplementary for "Secondary deficiency of neuraminidase 1 contributes to CNS pathology in neurological mucopolysaccharidoses via hypersialylation of brain glycoproteins"

### Supplementary materials

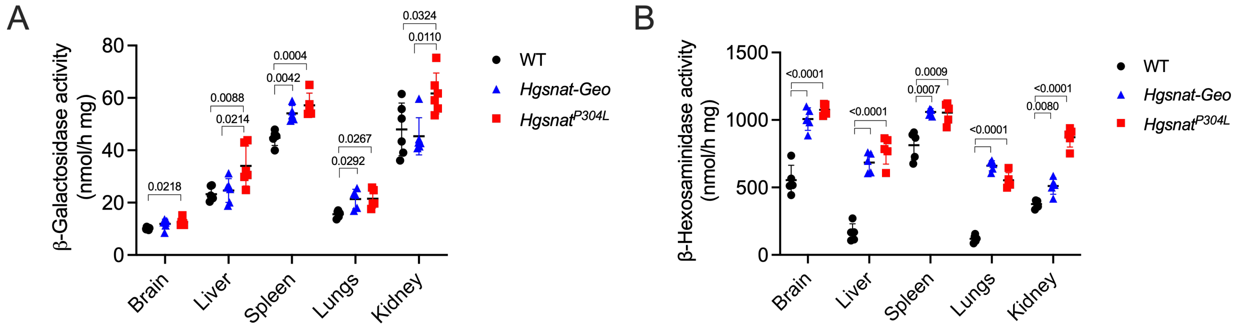


**Figure S1. Increased lysosomal biogenesis in tissues of MPS IIIC mouse models.**

Specific activities of β-galactosidase **(A)** and total β-hexosaminidase **(B)** were measured using the fluorogenic substrates, 4-methylumbelliferone-β-D-galactoside and 4-methylumbelliferyl-β-D-glucosaminide, respectively. The activities of both enzymes in tissues *Hgsnat-Geo* and *Hgsnat*$^{P304L}$ mice are increased or show a trend for an increase compared to WT counterparts. Individual data and means ±SD (n=5) are shown. P values were calculated by one-way ANOVA with a Tukey post hoc test.

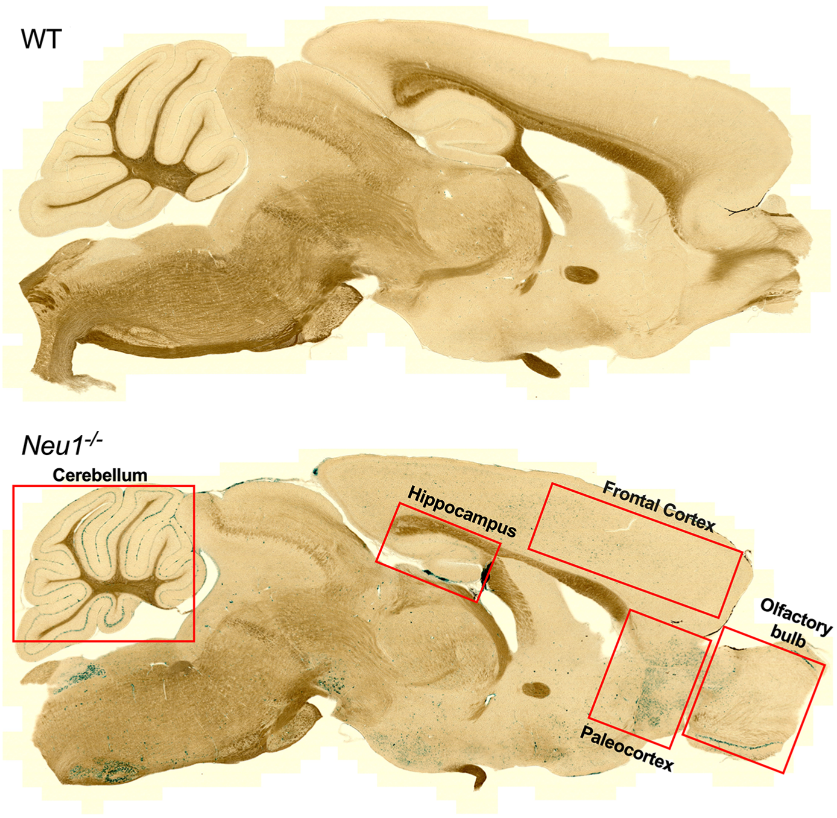


**Figure S2. X-GAL staining of mouse brain sections.**

Sagittal brain sections from 2-month-old WT and *Neu1*$^{-/-}$ mice were stained with the 5-bromo-4-chloro-3-indolyl-ß-D-galactopyranoside (X-GAL) substrate (blue) to visualise areas with high *Neu1* expression. The red boxes indicate regions that were dissected in WT, *Hgsnat*$^{304L}$ and *Hgsnat-Geo* mice for NEU1 enzyme activity assays. Images were taken at 10x magnification using an AxioScan instrument.

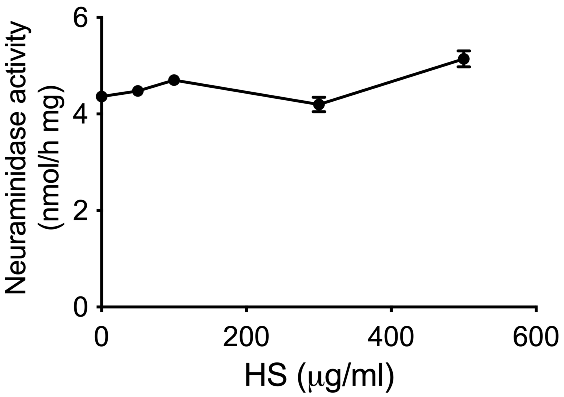


**Figure S3. Treatment of murine kidney homogenates with exogenous HS oligomers shows an absence of direct inhibition.**

Kidney homogenate of a WT mouse was treated with 0, 50, 100, 300 and 500 μg/ml of HS oligomers before measuring a total neuraminidase activity at pH 4.75. The total neuraminidase activity is not changed by HS. Graph shows means and SD of two technical replicas.

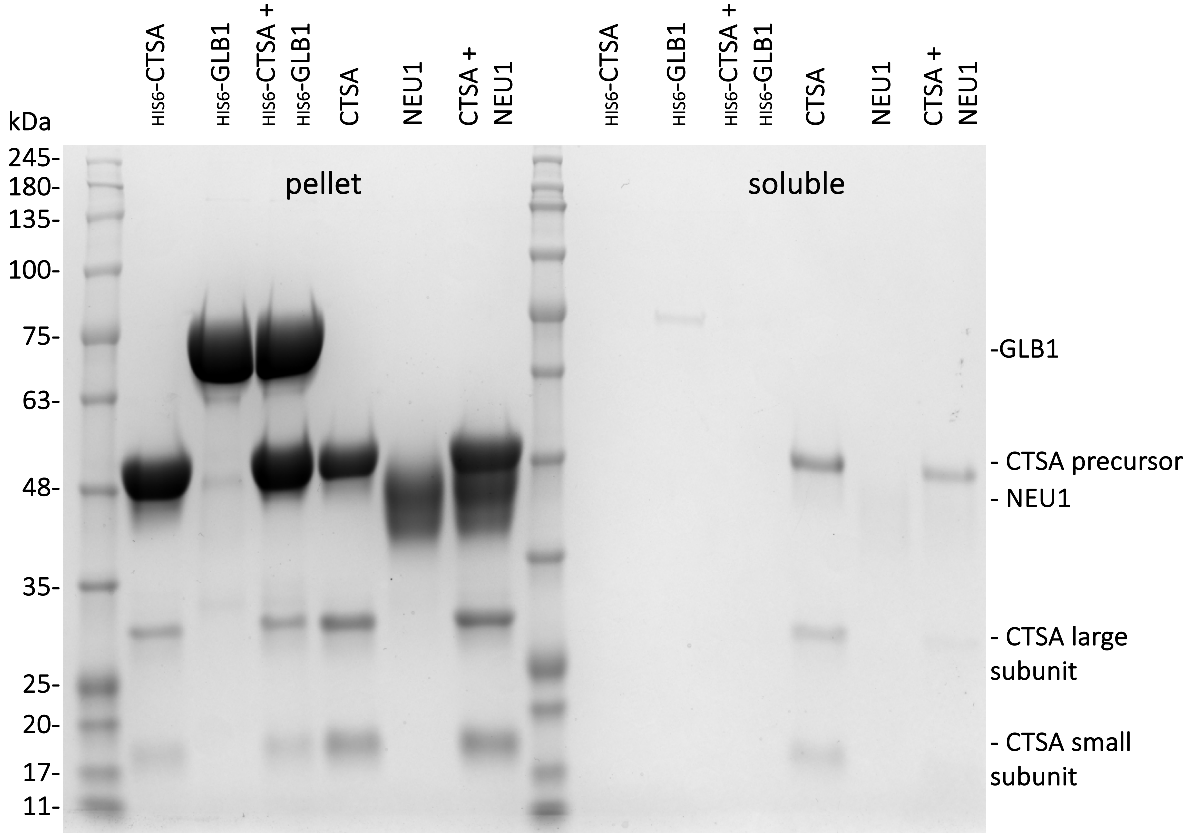


**Figure S4. HS causes precipitation of NEU1, CTSA and GLB1 *in vitro*.**

Purified recombinant NEU1, CTSA and GLB1 were incubated in various combinations in presence of 1 mg/mL of HS at pH 4.5. Equal volumes of soluble and precipitated fractions were analyzed by SDS-PAGE.

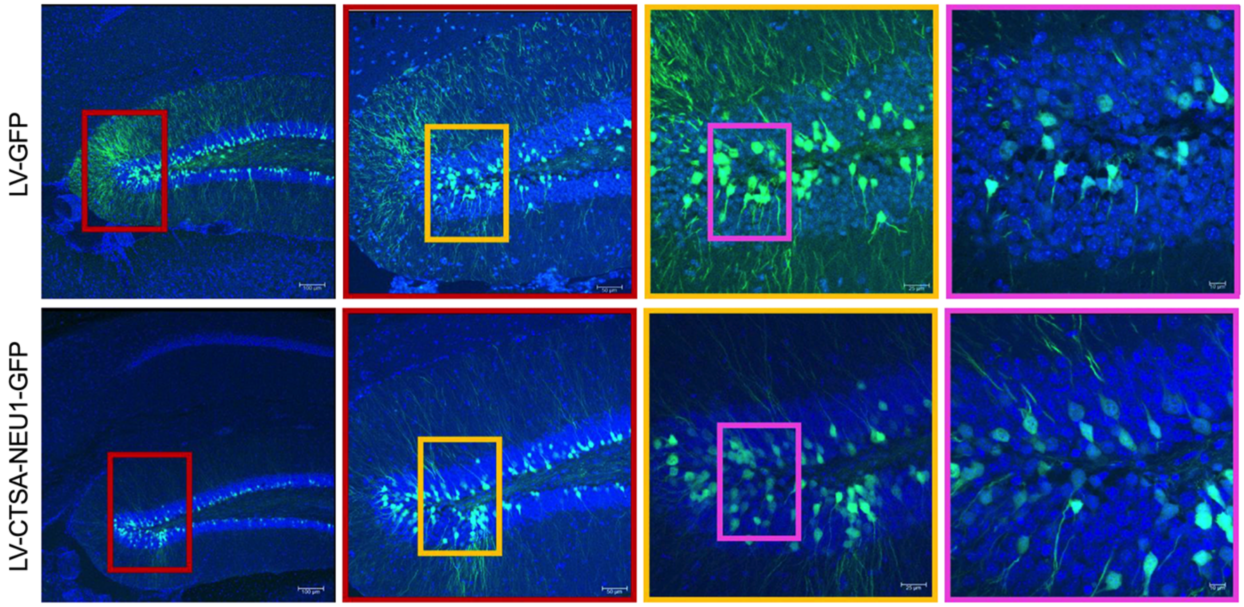


**Figure S5. Expression of GFP and NEU1-GFP proteins in hippocampus and dentate gyrus of mice treated with stereotaxic injections of LV-GFP and LV-CTSA-IRES-NEU1-GFP.**

GFP and NEU1-GFP proteins show high levels of expression in the neurons of hippocampus and dentate gyrus of mice six months after stereotaxic injections of LV-GFP and LV-CTSA-IRES-NEU1-GFP, respectively. Panels show representative confocal fluorescent images of hippocampus captured at different magnification. GFP-mediated fluorescence is shown in green. Nuclei were counterstained with DAPI (blue). Boxes show positions of zoomed areas. Scale bars: 100 µm and 50, 25 and 10 µm in zoomed images.

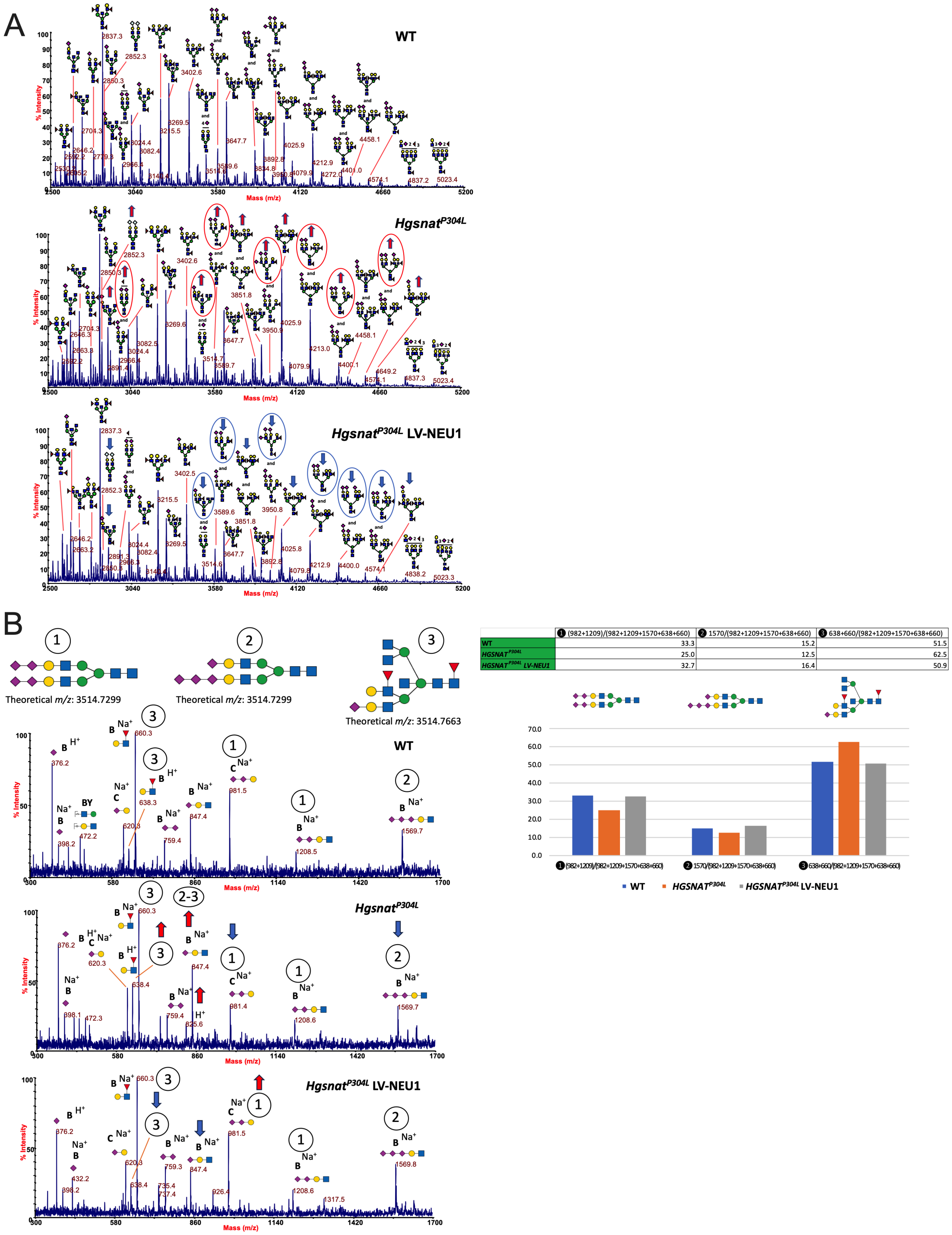


| | ❶ (982+1209)/(982+1209+1570+638+660) | ❷ 1570/(982+1209+1570+638+660) | ❸ 638+660/(982+1209+1570+638+660) |
|---|---|---|---|
| WT | 33.3 | 15.2 | 51.5 |
| *HGSNAT*$^{P304L}$ | 25.0 | 12.5 | 62.5 |
| *HGSNAT*$^{P304L}$ *LV-NEU1* | 32.7 | 16.4 | 50.9 |

**Figure S6. MALDI-TOF mass spectra show restored sialylation of hippocampal proteins in *Hgsnat*$^{P304L}$ mice treated with LV-CTSA-IRES-NEU1-GFP**

**(A)** Sialylation of N-linked glycans of hippocampal glycoproteins from 6-month-old WT, *Hgsnat*$^{P304L}$ and *Hgsnat*$^{P304L}$ mice treated with LV-CTSA-IRES-NEU1-GFP (LV-NEU1) was analyzed by MALDI-TOF MS.

Mass-spectra in the range between *m/z* 2500 to 5200 show increased levels of sialylated glycan species for untreated *Hgsnat*$^{P304L}$ compared to WT mice (red arrows) and reduced levels of sialylated glycans for LV-CTSA-IRES-NEU1-GFP treated mice compared to untreated *Hgsnat*$^{P304L}$ mice (blue arrows). Red circles mark the sialylated glyco-isomers that are enhanced in *Hgsnat*$^{P304L}$ mice and blue circles the ones reduced after the treatment according to the MS/MS analysis.

**(B)** Representative MS/MS analysis of the peak at *m/z* 3514.7 (low mass-range, between *m/z* 300 and 1700) shows that this species represent a mixture of three unique glycans with different positions of NeuAc moieties. Comparison of the relative intensity of the ions from non-reducing terminals revealed that fragments containing NeuAc 2,3 or 2,6-linked to Gal are increased in untreated and reduced in LV-CTSA-IRES-NEU1-GFP treated *Hgsnat*$^{P304L}$ compared to WT mice, while the intensity of fragment ions with 2,8-linked NeuAc dimers or trimers follows the opposite trend; they reduced in untreated *Hgsnat*$^{P304L}$ and return to WT levels in treated mice. This is reflected in the relative abundance of the three isomers at *m/z* 3514.7, as shown in the bar graph.

GlcNAc, blue square; Man, green circle; Gal, yellow circle; Neu5Ac, purple diamond; Neu5Gc, light blue diamond; Fuc, red triangle.

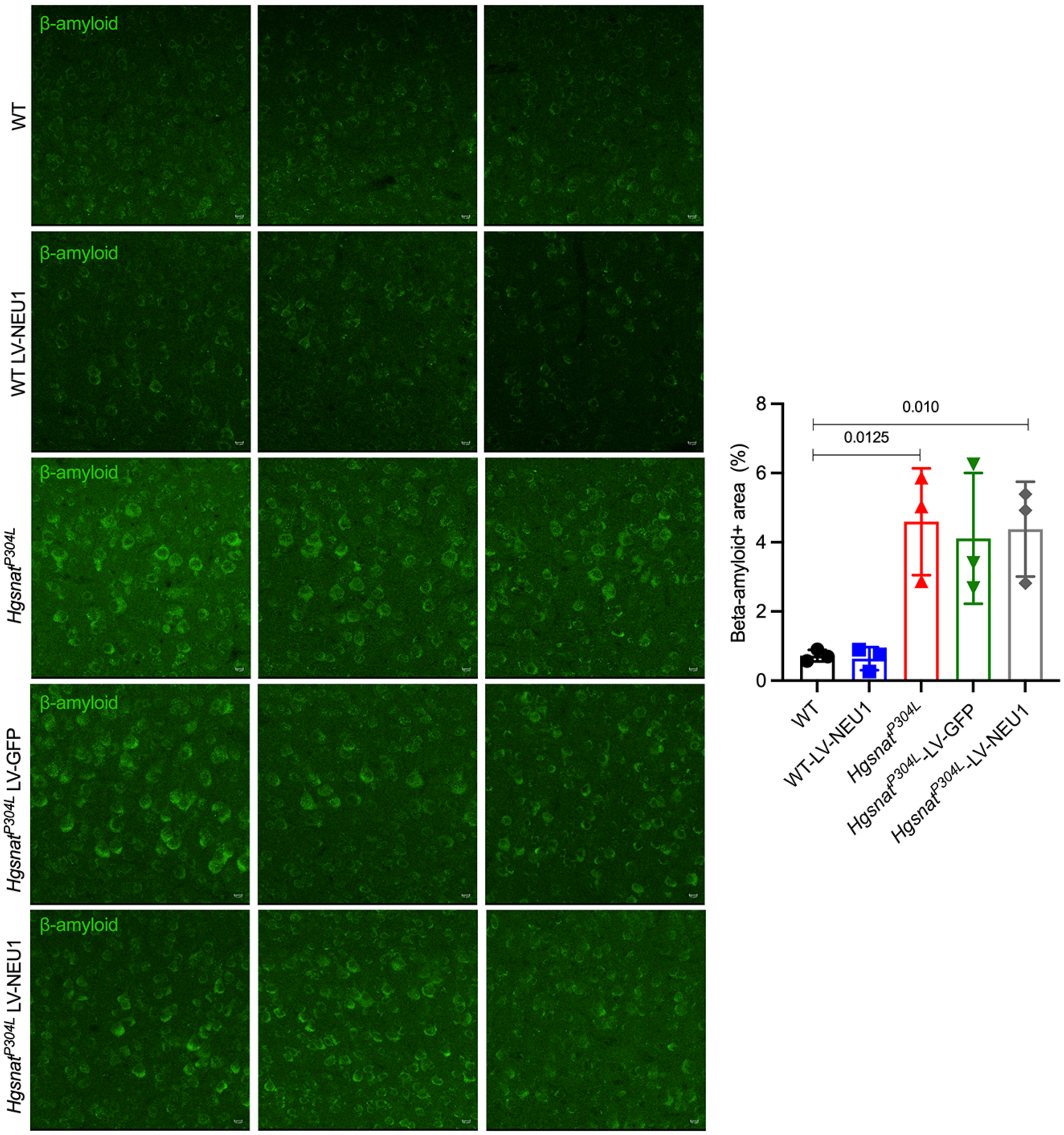


**Figure S7. *Hgsnat*$^{P304L}$ mice, treated with LV-CTSA-IRES-NEU1-GFP, show amyloid protein accumulation in level 4 and 5 pyramidal cortical neurons similar to that in *Hgsnat*$^{P304L}$ and sham-treated *Hgsnat*$^{P304L}$ mice.**

Representative confocal images of the somatosensory cortex of **(A)** WT mice, **(B)** WT mice injected with LV-CTSA-IRES-NEU1-GFP (LV-NEU1), **(C)** *Hgsnat*$^{P304L}$ mice, **(D)** *Hgsnat*$^{P304L}$ mice injected with LV-GFP, and **(E)** *Hgsnat*$^{P304L}$ mice injected with LV-CTSA-IRES-NEU1-GFP. Brain sections were stained with antibodies against beta-amyloid protein (green). Confocal images were taken using a 40x objective, scale bar: 25 µm. Data show means ± SD, n=5. Significance was determined by unpaired t-test.

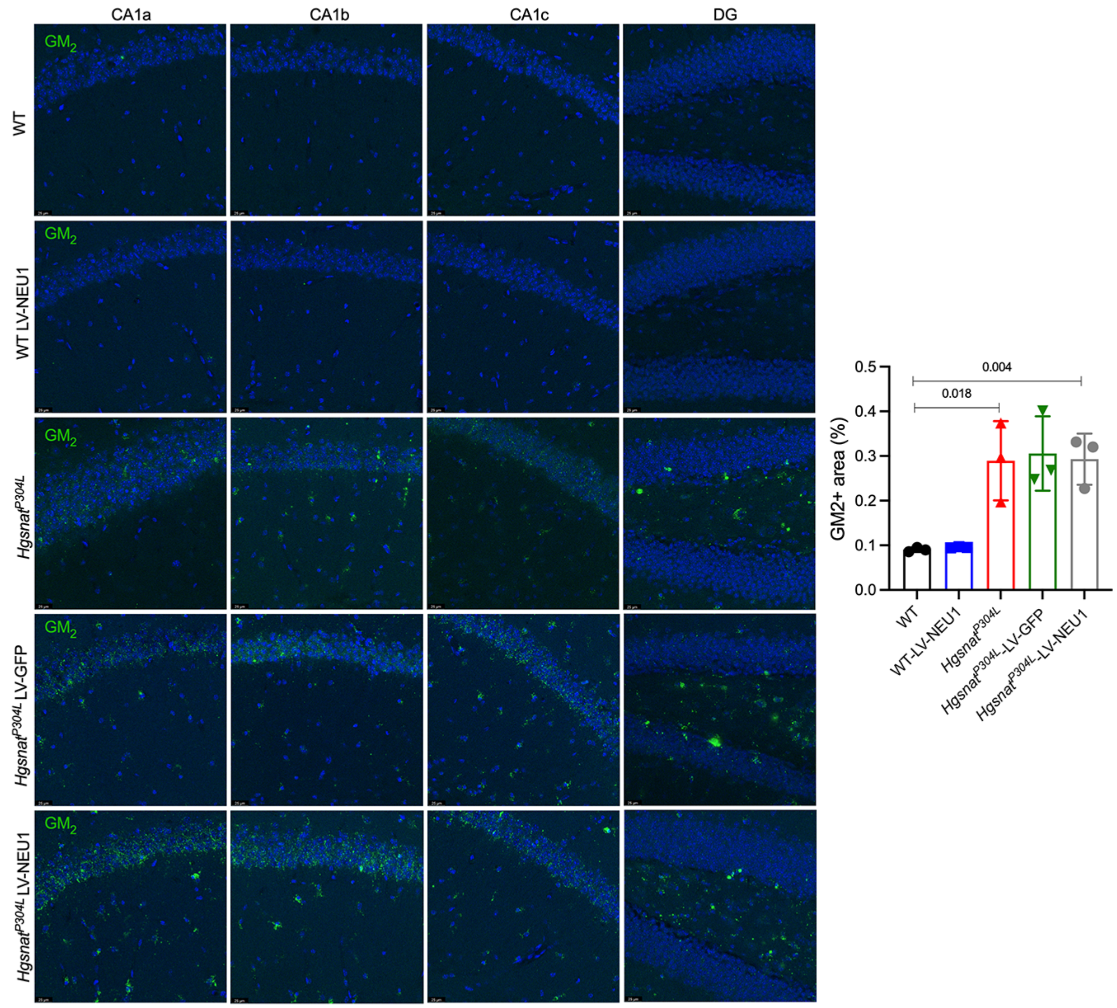


**Figure S8. *Hgsnat*$^{P304L}$ mice treated with LV-CTSA-IRES-NEU1-GFP show GM2 ganglioside accumulation in pyramidal hippocampal neurons similar to that in *Hgsnat*$^{P304L}$ and sham-treated *Hgsnat*$^{P304L}$ mice.**

Representative confocal images of four different regions of the hippocampus of **(A)** WT mice, **(B)** WT mice injected with LV-CTSA-IRES-NEU1-GFP (LV-NEU1), **(C)** *Hgsnat*$^{P304L}$ mice, **(D)** *Hgsnat*$^{P304L}$ mice injected with LV-GFP, and **(E)** *Hgsnat*$^{P304L}$ mice injected with LV-CTSA-IRES-NEU1-GFP. Brain sections were stained with antibody against GM2 ganglioside (green). The nuclei were counterstained with DAPI (blue). Confocal images were taken using a 40x objective, scale bar: 25 µm. Data show means (± SD), n=3. Significance was determined by unpaired t-test.

**Supplementary Table S1. Human MPS patients and non-MPS controls used in the study**

| Identification | Disorder | Cause of Death | Age | Sex | Ethnicity |
|---|---|---|---|---|---|
| 662 | Control | Accident, multiple injuries | 12 | Female | White |
| 754 | Control | Asthma | 11 | Female | Native Hawaiian or Other Pacific Islander |
| 1266 | Control | ASCVD (Arteriosclerotic Cardiovascular Disease) | 42 | Male | White |
| 4641 | Control | Asthma | 24 | Female | Black or African-American |
| 5287 | Control | Accident, multiple injuries | 23 | Female | White |
| 5813 | Control | ASCVD (Arteriosclerotic Cardiovascular Disease) | 20 | Male | Black or African-American |
| 5977 | Control | Smoke inhalation | 6 | Female | White |
| 561 | MPS I | Complications of disorder | 6 | Female | White |
| 902 | MPS II | Complications of disorder | 42 | Male | White |
| HBCB_18_01_OC | MPS II | Unknown | 13 | Male | White |
| -3617 | MPS IIIA | Complications of disorder | 12 | Female | White |
| 563 | MPS IIIA | Complications of disorder | 11 | Female | White |
| 6194 | MPS IIIC | Complications of disorder | 20 | Male | Black or African-American |
| 5411 | MPS IIID | Complications of disorder | 24 | Female | White |
| 5424 | MPS IIID | Complications of disorder | 23 | Female | White |
